# Organic Components Modulate the Morphology of Respirable Aerovirology-Relevant Aerosols

**DOI:** 10.64898/2026.01.16.699978

**Authors:** Deepak Sapkota, Yuhui Guo, Amrita Chakraborty, James Hu, Harris Xie, Ian Wu, Moon J. Kim, Hui Ouyang

## Abstract

Airborne transmission of pathogens occurs via aerosol particles, whose morphology provides insights into the microenvironments that pathogens experience. Aerosol morphology includes particle size, shape, phase state, and chemical homogeneity, yet systematic studies remain limited. Here, we characterized model bioaerosol morphologies generated from (1) NaCl–organic two-component mixtures, (2) common cell culture media, and (3) artificial respiratory fluids. Particles were collected using a virtual impactor and Andersen cascade impactor and analyzed by scanning electron microscopy (SEM) and energy-dispersive X-ray spectroscopy (EDX). Results show that organic components modulate the morphology: dipalmitoylphosphatidylcholine (DPPC) promotes organic-inorganic phase separation while proteins prohibit formation of large crystals and leads to better mixing among components. At 30% RH with a drying period of 10 seconds, most aerosols appeared desiccated, though NaCl-glucose, DMEM-complete-media and artificial saliva with mucin remained semi-solid or gel-like. Among all formulations examined EMEM-complete-media and artificial saliva (non mucin) show a size-dependent morphology. Our study demonstrates how chemical composition and size alters surrogate bioaerosol phase (semi-solid or solid) and morphology and provides new insights into the microenvironment of aerosol particles for aerovirology investigations.

## 1. Introduction

Laboratory-controlled aerovirology studies typically involve the generation, evaporation, suspension, and sampling of virus-laden bioaerosols, followed by gene copy and viability analysis (Groth et al. 2024; Ma et al. 2021; Haddrell and Thomas 2017). These studies effectively simulate airborne virus transmission and offer a robust framework for systematically investigating how environmental conditions impact the airborne pathogen transmission, particularly during the suspension stage, such as airflow and ventilation (Thornton et al. 2022), relative humidity (RH) (Motos et al. 2024; Ahlawat et al. 2020; Kormuth et al. 2018), temperature (Prussin et al. 2018; Hermann et al. 2007; Lowen et al. 2007), and UV exposure (Xia et al. 2022; Qiao et al. 2021; Raeiszadeh and Adeli 2020; Walker and Ko 2007; Tseng and Li 2005; Jensen 1964). After bioaerosol droplets undergo rapid evaporation within seconds (Walker et al. 2021; Wang et al. 2021; Yang and Marr 2011; Xie et al. 2007; Nicas et al. 2005), bioaerosol particles can remain suspended for minutes to hours (Eilts et al. 2021; Sattar et al. 2016; Thatcher et al. 2002) after reaching equilibrium, contributing to both short- and far-field transmission (Dong et al. 2024; Jiang et al. 2024; Xu et al. 2024; Chien et al. 2022; Hsu et al. 2022; Leung 2021). While prior studies have characterized particle size distribution and biological properties such as gene copy number and infectivity, a mechanistic understanding of virus inactivation during the suspension stage remains limited. Specifically, improved understanding of microenvironmental conditions within particles is needed to uncover the underlying processes governing viral fate.

The morphology and organic-inorganic phase separation of virus-free surrogate bioaerosols, governed primarily by their organic and inorganic constituents rather than the presence of virus itself, is a critical physicochemical property that provides insight into the microenvironmental conditions surrounding pathogens in bioaerosol particles. Interestingly, the concept of morphology in this context extends beyond the traditional definition, which typically refers to structural shape (e.g., spherical, cubic, or irregular aggregates). It also includes aspects such as chemical homogeneity, the spatial distribution of components, and phase characteristics (e.g., liquid, solid, semi-solid or glassy states). While numerous studies have explored the morphology of atmospheric aerosols (Lei et al. 2025; Yoo et al. 2024; Fu et al. 2020; Shiraiwa et al. 2013; Freedman et al. 2010; Moffet et al. 2010; McDonald and Biswas 2004), surrogate bioaerosols generated in aerovirological studies differ significantly in composition, containing complex mixtures of inorganic salts, proteins, sugars, lipids, and other organics. These constituents directly influence particle morphology upon drying. Morphological features, such as core-shell structures, can lead to the phase separation of organics from salts, potentially shielding embedded viruses from harsh chemical environments (Pan et al. 2025). As such, morphology may serve as a valuable property for identifying the physicochemical factors that impact virus transmission and can improve the comparability of findings across aerovirological studies.

Importantly, the role of morphology becomes even more critical under low RH conditions (<50%), where water evaporates and particles become dry nuclei. These conditions are common in indoor environments, particularly during winter in temperate climates (Jones et al. 2022; Tamerius et al. 2013; Lowen et al. 2007). Some viruses, including respiratory syncytial virus (RSV), human rhinovirus (HRV), MS2, □6, Middle East respiratory syndrome coronavirus (MERS-CoV), severe acute respiratory syndrome coronavirus (SARS-CoV), and influenza viruses, demonstrate enhanced survival rates at low relative humidity (RH), though the underlying mechanisms remain unclear (Niazi et al. 2021, 2023; Lin and Marr 2020; Prussin et al. 2018; van Doremalen et al. 2013; Chan et al. 2011; Lowen et al. 2007). However, the morphological state of surrogate bioaerosols dried at low RH conditions has received limited attention in aerovirology. Only a few recent studies have examined post-aerosolization morphology using electron microscopy on selected model solutions, such as Dulbecco’s Modified Eagle Medium (DMEM) (Alexander et al. 2022), Eagle’s Minimum Essential Medium (EMEM) (Oswin et al. 2022), artificial saliva (AS), and deep lung fluid (Meng et al. 2025; Tian et al. 2024; Walker et al. 2021), Phosphate-Buffered Saline (PBS) and Luria-Bertani (LB) broth (Otero-Fernandez et al. 2024), and porcine respiratory fluid (Groth et al. 2023). From these studies, phase status depends on the RH value greatly and, for dry particles, mucin seems to delay crystallization for inorganic salts and produce compact particles with smooth surface. Yet, a systematic morphological investigation across commonly used laboratory aerosolization solutions and quantitative chemical distribution are still lacking.

Previous studies of respiratory bioaerosols have primarily focused on submicron particles (Groth et al. 2022, 2023) or coarse particles ≥5 µm (Otero-Fernandez et al. 2024; Tian et al. 2024), leaving the intermediate size range of 1–5 µm comparatively underexplored. This size range is highly relevant for airborne disease transmission, as particles smaller than 5 µm are respirable, can penetrate into the lower respiratory tract (Sadeghi et al. 2025; Wang et al. 2021; Milton et al. 2013), and may directly contribute to lower respiratory infections (Verwey and Nunes 2020). Particle size is closely linked to aerosol generation mechanisms and anatomical origin within the respiratory tract. During respiratory activities such as speaking and coughing, aerosols generated in the bronchioles typically have mean diameters near 1 µm, whereas particles originating from the larynx average around 5 µm (Morawska et al. 2022; Johnson et al. 2011) Because respiratory fluid composition varies by generation site, aerosols are expected to exhibit coupled size–composition dependencies (Prussin et al. 2023). Bronchiolar fluids are enriched in pulmonary surfactants (Kakeshpour et al. 2025), whereas laryngeal secretions contain notable concentrations of mucins (Fahy and Dickey 2010). (Santarpia et al. 2024) provide a detailed review of the pathways of virus□laden aerosols generated during respiratory activities, emphasizing the co□variation of particle size and chemical composition across different regions of the respiratory tract. These size□ and composition□dependent differences have direct implications for aerosol morphology and pathogen persistence. Mucin□rich particles promote early□stage phase separation, including core–shell and inclusion structures, and increasing mucin concentrations are associated with reduced viral inactivation (Alexander et al. 2022). Together, these findings underscore the importance of simultaneously considering particle size and chemical composition when investigating respiratory bioaerosol morphology and pathogen inactivation during airborne transmission.

In this work, we systematically characterize the morphologies of virus-free surrogate bioaerosols at low RH within the size range of 1-5 µm, derived from (1) NaCl–organic two-component systems, (2) common cell culture media, and (3) widely used artificial respiratory model fluids. Using SEM and EDX analysis of particles collected with an Andersen impactor, we show how chemical composition governs phase separation between organic and inorganic components at both global and local scales. Specifically, NaCl–glucose aerosols form core–shell structures, while mucin and BSA promote well-mixed states. In contrast, DPPC induces pronounced phase separation. Cell culture media produce globally well-mixed particles (with localized separation upon addition of serum), whereas for the selected model respiratory fluids the addition of mucin to artificial saliva or DPPC to artificial lung fluid distinctly alters phase state and morphology. Furthermore, under identical drying conditions, the phase state of surrogate bioaerosols (semi-solid versus solid) depended on both chemical composition and particle size, with serum shifting larger particles from solid to semi-solid states. By coupling structural imaging with compositional mapping, this study provides new insights into the physicochemical states adopted by bioaerosols during suspension, offering a framework for understanding the microenvironments in which pathogens reside and the mechanisms of airborne virus transmission.

## 2. Methods

### 2.1 Solution preparation

We prepared three types of suspension solutions: two-component systems containing sodium chloride (NaCl) and organics, cell culture media, and model respiratory fluids. For two-component NaCl-organic system, we selected glucose (Thermo Scientific Chemicals, AAA1682836), bovine serum albumin (BSA, Fisher Scientific, BP9703100), porcine stomach mucin (Millipore Sigma, M1778,Type III, bound sialic acid 0.5-1.5%, partially purified powder), and DPPC (Avanti Polar Lipids, AL, USA) as representative sugars, proteins, and lipids found in respiratory fluids. We maintained the NaCl concentration at 6 g/L, consistent with the levels in artificial lung fluids (ALF), Eagle’s Minimal Essential Medium (EMEM), DMEM, and other in vitro lung bioaccessibility fluids (Boisa et al. 2014; Moss 1979). We also adjusted the organic concentration to 6 g/L to achieve an organic-to-inorganic mass ratio of 1 in the two-component NaCl-organic systems.

For the cell culture media, we purchased DMEM, EMEM, and PBS (all from Corning, NY, USA) and used them without modification. To create complete cell culture media, we added 10% (v/v) Fetal Bovine Serum (FBS) (Fisher Scientific, PA, USA) and 1% (v/v) Penicillin-Streptomycin-L-glutamine (Corning, NY, USA) to DMEM and EMEM.

We also formulated model respiratory fluids. We obtained artificial saliva (AS-P, 1700-0304) from Pickering Laboratories (Mountain View, CA, USA) and used it without modification. In the lab, we prepared an alternative artificial saliva (AS-W) following Woo et al. (Woo et al. 2010), and we also produced artificial saliva without mucin (AS-NM) using the same method. We purchased ALF from Pickering Lab (Cat. 1700-0808, Mountain View, CA, USA) and used it as received. To mimic deep lung fluid composition (Hassoun et al. 2018), we added 48 mg of DPPC to 10 ml of ALF along with 30 ml ethanol (Fisher Scientific, Cat: 64-17-5, 99.5% Purity), producing a modified lung fluid (ALF-DPPC). Supplemental Information lists detailed chemical compositions for each solution (**Table S1–S10**).

### 2.2 Aerosol generation and characterization

To generate aerosol particles, we used a Single-Jet Blaustein Atomizing Module (CH Technologies, NJ, USA) alongside a syringe pump (Braintree Scientific, Inc., MA, USA). We positioned the atomizer nozzle directly inside a stainless-steel tube (1.4-inch inner diameter, McMaster sanitary fittings) with a total length of 44 inches (**Figure S1**). The syringe pump fed the suspension solution into the nozzle at a rate of 0.15 mL/min, while we supplied compressed air at 30 psi. We introduced additional dilution air at the tube inlet, which resulted in a combined flow rate of 31 L min^-1^ at the aerosol generator, as measured by a flow meter (model 4040, TSI MN, USA). We monitored relative humidity and temperature at four locations along the tube using a probe (Ahlborn, Germany) and recorded the relative humidity in the sanitary tube as 27.8 ± 3.0%. To neutralize excess electrostatic charges on aerosol particles, we affixed a Po-210 strip to the inner wall near the tube exit. We controlled the downstream flow to allow the desired aerosol flow rate to exit the tube, with excess flow escaping through a vent located near the aerosol generator. We conducted experiments using a 5 L min^-1^ downstream flow rate to maintain a consistent drying time of 10 seconds.

To separate particles by size, we placed a custom-built virtual impactor (VI) at the exit of the generation tube: smaller particles followed the major flow, while larger particles concentrated into the minor flow (Li et al. 2025; Eilts et al. 2023; Loo and Cork 1988; Marple and Chien 1980). We controlled a central air stream with a mass flow controller and incorporated it to minimize small-particle contamination in the minor flow line (Li and Lundgren 1997; Chein and Lundgren 1993). The virtual impactor provided a cutoff size of 2 µm, with corresponding flow rates of 9.7 L min^-1^ in the major flow and 0.3 L min^-1^ in the minor flow. We achieved flow control with a mass flow controller (MFC) (Aalborg Instruments and Controls, Inc., USA), using either a vacuum pump (Gast Manufacturing Inc MI, USA) or a compressed air line. We introduced larger particles from the minor flow into an eight-stage non-viable Andersen Cascade Impactor (ACI) (Tisch Environmental, OH, USA), collecting them on silicon wafers (Ted Pella, Inc., USA) placed on different stages. We added dilution air to maintain a total flow of 28.3 L min^-1^, which is the standard operational flow rate of the ACI (Andersen 1958) to ensure particles with aerodynamic diameters of 0.7–5.8 µm deposited across multiple stages (stages 2–6). We monitored and recorded the exit flow relative humidity of the ACI as 2.8 ± 0.2%. We optimized the collection period, ranging from two to ten minutes, to minimize particles stacking at each collection spot, especially on stage 4. We measured aerosol particle size distributions using an Aerodynamic Particle Sizer (APS, model 3321, TSI, MN) and a Scanning Mobility Particle Sizer (SMPS, model S3938L56, TSI, MN).

### 2.3 Aerosol imaging

After collection, we carefully removed the silicon wafers from the ACI and transported them to the core facility for SEM imaging (Zeiss Sigma 500 VP). We used secondary electron mode for higher resolution and surface detail, and an energy-dispersive X-ray (EDX) spectrometer to perform elemental analysis. For each sample, we optimized the working distance and accelerating voltage, ranging from 3 mm to 10 mm and 0.6 kV to 2 kV, respectively. We optimized the acceleration voltage to exclude imaging artifacts from electron-beam heating, even enabling the capture of well-defined solid spheres from particles generated using a 20 g L□¹ glucose solution (**Figure S1**). During EDX elemental analysis, we conducted a line scan using a working distance of 14–16 mm and an accelerating voltage of 20 kV. We selected a dwell time of 800 ms per point for the line scan, with an overall scan duration of approximately 14 minutes per particle. We set the electron beam aperture to 120 µm during EDX analysis. We report the normalized element intensity as a function of location along the scan.

### 2.4 Colocalization analysis for elements

To quantify elemental colocalization, we performed correlation analysis by calculating Pearson correlation coefficients (ρ) for EDX line scan data and use ρ ≥ 0.7 as evidence for strong co-localization, consistent with prior literature (Helfferich et al. 2025; Schober and Schwarte 2018) We calculated and reported the average and standard error for the correlation coefficient across multiple particle samples (n≥3).

## 3. Results

### 3.1 Organic composition governs phase separation (NaCl-glucose, NaCl-DPPC) or mixing (NaCl-mucin, NaCl-BSA) in NaCl–organic bioaerosols

First, we confirmed the generation of sub–5 µm particles (**Figure S2**). We used APS and SMPS which report particle size in aerodynamic diameter and mobility diameter separately. Here we present these results as it is without converting between metrics for clarity. Our setup reliably generated a bimodal distribution in number concentration, with modes near ∼60 nm and 1 µm. By employing the virtual impactor, we filtered out nanoparticles, resulting in aerosol number and volume distribution peaks at approximately 2 µm and 3 µm, respectively. Particles are collected across Andersen Cascade Impactor (ACI) stages 2 through 6, covering aerodynamic diameter ranges from 0.7 to 5.8 µm (**Table S11**). We validated these ACI cutoff sizes using 2 µm and 3.1 µm polystyrene latex particles (**Figure S3**).

Next, we investigated aerosol morphologies in two-component NaCl–organic particles consisting of different organic components with a fixed 1:1 organic-to-inorganic mass ratio (6 g/L each) solutions. We selected glucose, BSA, mucin, and DPPC for the organic component to represent sugars, proteins, mucus, and lipids, respectively. Our combined SEM and EDX analyses reveal distinct 3D structures and chemical distributions in NaCl–organic aerosols, with morphology strongly influenced by the type of organic component (**Figure 1, S4-S7**). First, none of the four systems display X-ray intensity profiles characteristic of uniform solid spheres as in BSA (**Figure S8** with parabolic elemental intensity profiles that peak at the center). Instead, dips in Na and Cl intensities suggest irregular internal structures (**Figure S9-S10**). For example, NaCl–glucose and NaCl–DPPC samples show gaps between larger NaCl crystals, while NaCl–mucin and NaCl–BSA particles exhibit surface dents or possible hollow structures. Second, NaCl–glucose and NaCl–DPPC mixtures exhibit strong phase separation from both SEM imaging as well as the near-zero correlation coefficient for Na-C (**Figure 1d & h**). In NaCl–glucose, NaCl forms distinct cubic cores surrounded by glucose, yielding a well-defined core–shell morphology (**Figure 1a&b, S4**), matching previous imaging observations (Lee et al. 2019; Ray et al. 2019). NaCl–DPPC also shows strong phase separation, with salt crystals either exposed at one end or encapsulated in the organic suggested by Na signals peak in regions lacking visible crystals (**Figure 1e&f, S5**). Due to DPPC’s low water solubility, two extreme morphologies were also observed: (1) pure NaCl crystalline particles and (2) smooth, irregular DPPC-rich particles (**Figure S5**). In contrast, NaCl–BSA and NaCl–mucin form more well-mixed homogeneous non-spherical particles without large, distinct salt crystals. Their high Na–C correlation coefficients (∼0.8), compared to NaCl–Glucose and NaCl–DPPC systems, indicate better mixing between inorganic and organic components. However, SEM images of NaCl-mucin reveal small salt crystals embedded within the matrix or protruding from the surface (**Figure S6**), suggesting that although bulk mixing is achieved, localized phase separation still occurs. Therefore, the organic phase composition determines the phase mixing state in two-component NaCl–organic crystals under the conditions we investigated.

**Figure 1.**
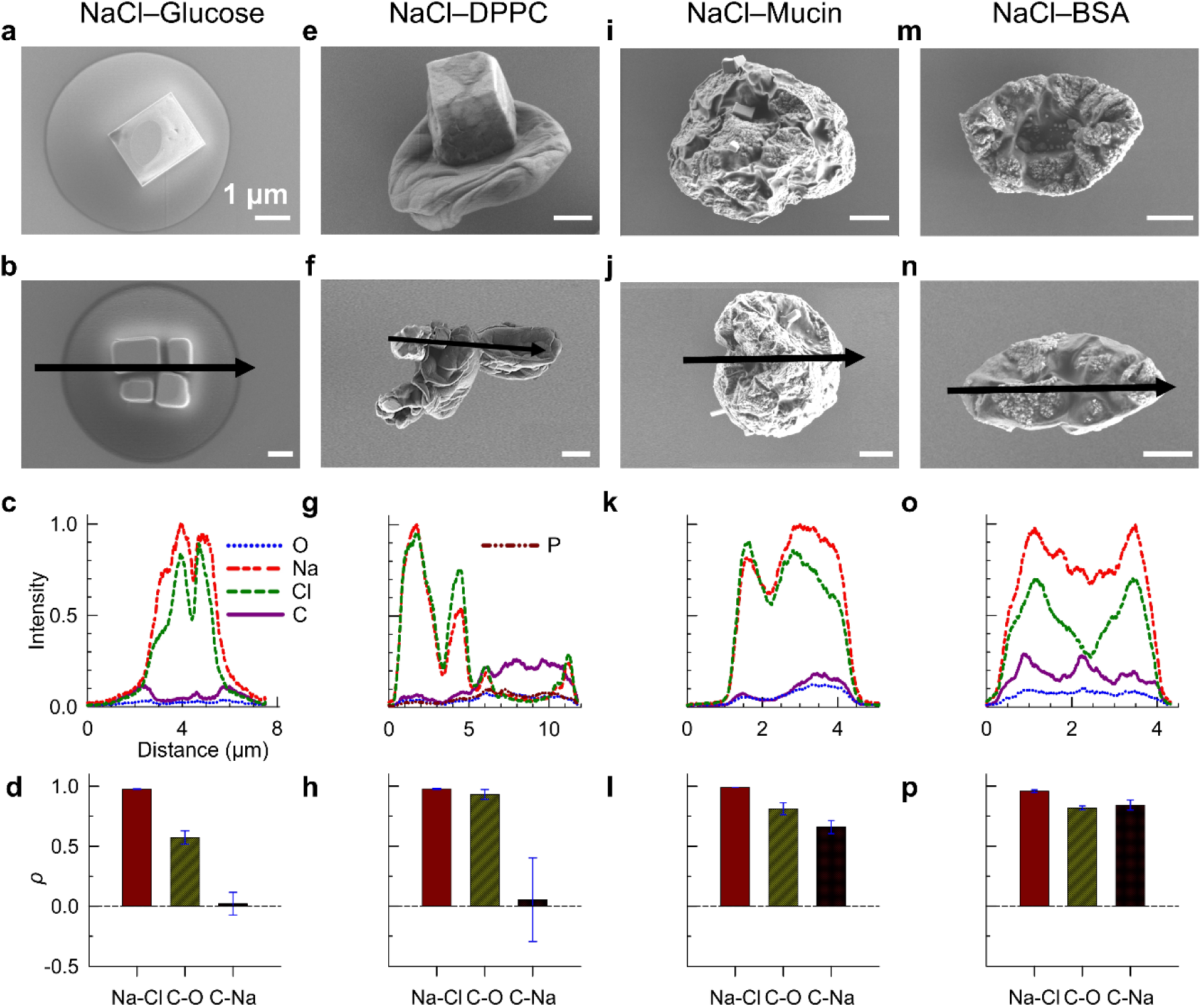
Organic composition governs phase separation (NaCl-glucose, NaCl-DPPC) or mixing (NaCl-mucin, NaCl-BSA) in NaCl–organic aerosol particles. Panels display representative SEM morphologies, EDX elemental distributions and Pearson correlation coefficients (ρ) between different element pairs. All samples were derived from two-component organic-inorganic mixtures (OIR = 1) and dried at ∼30% relative humidity for 10 seconds.

### 3.2 Common cell culture media lead to globally chemically well-mixed aerosols and addition of serum leads to localized phase separation

Next, we imaged virus-free surrogate bioaerosols particles from commonly used cell culture media. The chemical compositions of the solutions are summarized in **Table S1-S5**. The chemical complexity of common cell culture media exerts a substantial influence on the morphology and elemental distribution of aerosolized particles (**Figure 2, S11-S15**), often resulting in a wide range of particle shapes and internal structures. SEM imaging and EDX elemental analysis reveal that PBS, a medium comprised exclusively of inorganic salts, forms single or multicrystalline particles with highly uniform elemental distributions across particles of varying sizes. The EDX maps consistently show Pearson correlation coefficients close to 1 between salt pairs, indicating well-mixed compositions; however, nonuniform intensity profiles (dips in intensity) suggest possible voids within multicrystalline particles.

**Figure 2.**
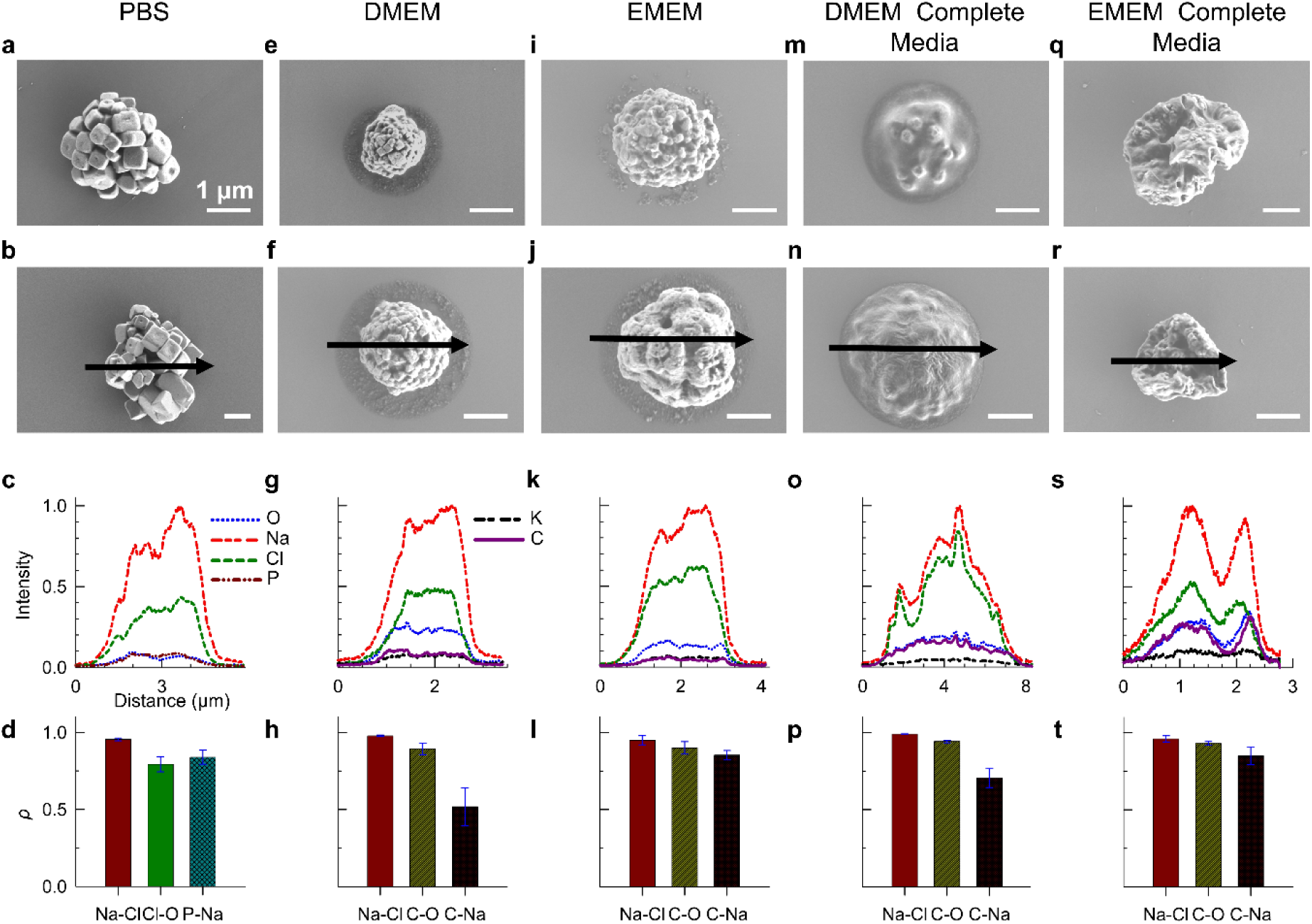
Common cell culture media leads to chemically globally well-mixed aerosol particles, whereas the addition of serum induces localized phase separation. The top two rows show representative particles produced from phosphate-buffered saline (PBS), Dulbecco’s Modified Eagle Medium (DMEM), Eagle’s Minimum Essential Medium (EMEM), DMEM complete media, and EMEM complete media. The bottom two rows display corresponding elemental intensity profiles and Pearson correlation coefficients. PBS particles exhibit crystalline morphology, while DMEM and EMEM produce near-spherical, well-mixed particles with small satellite droplets and minimal organic–inorganic separation. In contrast, media supplemented with fetal bovine serum (DMEM/EMEM + FBS) generate particles containing encapsulated salt crystals surrounded by an organic matrix, with localized phase separation, with satellites notably absent.

DMEM and EMEM are complex mixtures containing inorganic salts as well as organic components such as glucose, amino acids, and vitamins, and, upon aerosolization, both media produce near-spherical particles with rough surfaces and small dents, and SEM imaging reveals no evident organic–inorganic phase separation. This observation is supported by elemental mapping, which demonstrates high Pearson correlation coefficients (ρ > 0.75) for Na–Cl, C–O, and C–Na pairs with the exception of moderate correlation coefficient (ρ = 0.52 ± 0.12) for C-Na pair in DMEM, indicating a well-mixed internal structure.

A distinctive morphological feature in DMEM and EMEM particles is the clustering of small satellite particles around a larger main particle, likely resulting from fragmentation of loosely bound organic regions during impaction or splashing of water-rich content followed by re-structuring during the drying process, consistent with previous report (Freedman et al. 2010).

When both DMEM and EMEM are supplemented with 10% (v/v) fetal bovine serum (FBS) and 1% penicillin (v/v), forming complete media, the particle morphology undergoes a distinct transformation in four aspects. Firstly, satellite structures seen in unsupplemented samples disappear entirely (**Figure 2m**, **2n**, **2q**, **2r, S14, & S15**). Secondly, we observed numerous small salt crystals encapsulated by organic-rich domains among DMEM complete media particles, suggesting localized phase separation. We identified small NaCl crystals by EDX spectra, with Na and Cl peaks localized at specific crystal regions (**Figure 2o**). The amorphous matrix, which likely originates from serum proteins, embeds or coats these crystals and encapsulates the small salt crystals within a thick organic layer (**Figure S16**). This localized phase separation is masked by high C–Na correlation coefficients (**Figure 2p**), indicating global co-localization of organic and inorganic components, though tiny salt crystals remain embedded within organic-rich domains. Thirdly, even though elements are well-mixed (high correlation coefficients for all pairs (**Figure 2t**), we observe complex surface structures for desiccated EMEM complete media particles with dents and concaves. Major elements (Na, Cl, C, and O) peak at the same location with two distinct peaks at the edge of a cave. Lastly, DMEM complete media particles show spreading behavior upon impaction similar to NaCl–glucose systems, with projected-area equivalent diameters larger than the expected particle diameters (**Figure S17**), indicative of a semi-solid or gel-like state resulting in partial flattening.

### 3.3 Organic composition (urea, mucin, and lipid) modulates the morphology and organic-inorganic separation in particles derived from model respiratory fluids

Lastly, we investigated aerosol particles generated from model respiratory fluids, including AS and ALF formulations, and examined the influence of organic composition including urea, mucin and lipid present in these fluids. AS-P contains urea as the primary organic component and multiple inorganic salts. AS-P aerosol particles display core–shell morphologies (**Figure 3a-c & S17**) with visible phase separation: crystalline salt cores, predominantly KCl due to the low Na:K ratio (0.1), surrounded by a coating. EDX analysis confirms a strong K–Cl co-localization (**Figure 3e**). AS-P particles also exhibited projected-area equivalent diameters larger than their expected particle diameters (**Figure S17**), consistent with a soft, gel-like structure and spreading behavior upon impaction. EDX analysis indicated that, aside from the distinct KCl crystals, the remaining elements appeared well-mixed with EDX peaks at the edges of the crystals. In contrast, artificial saliva without mucin (AS-NM), which has an extremely low organic-to-inorganic mass ratio (0.04), produced dry, near-spherical particles with prominent crystalline features (**Figure 3f-h & S19**). EDX spectra confirmed these crystals as KCl, showing distinct K and Cl peaks, while other elements appeared broadly mixed but spatially offset from the KCl signals, suggesting microphase separation (**Figure 3h-j**). Adding porcine stomach mucin (3 g/L) to artificial saliva following Woo.et al’s formula (AS-W) substantially altered particle morphology and phase state (**Figure 3k-m & S20**). At the current RH (∼30%) and drying conditions (∼10 s drying time), AS-W particles appeared semi-solid, spreading upon impaction to form circular stain rings, similar to the gel-like phase of NaCl-glucose. EDX analysis showed a relatively higher Pearson coefficients among different element pairs confirming globally homogeneous mixing (**Figure 3o**). However, the offset of K and Na peaks suggests local phase separation for K-rich and Na-rich components (**Figure 3m-n)** and multiple peaks of Na-, C-and O- shows at the same locations suggesting that NaHCO_3_ might be localized (**Figure 3n**). While small crystals were locally observed, mucin inhibited the formation of large salt crystals, promoting smoother morphologies and more spherical particle structures. These findings demonstrate that mucin plays a critical role in enhancing phase mixing, maintaining semi-solid states, and suppressing crystallization, leading to particle morphologies distinct from those of saliva formulations lacking mucin.

**Figure 3.**
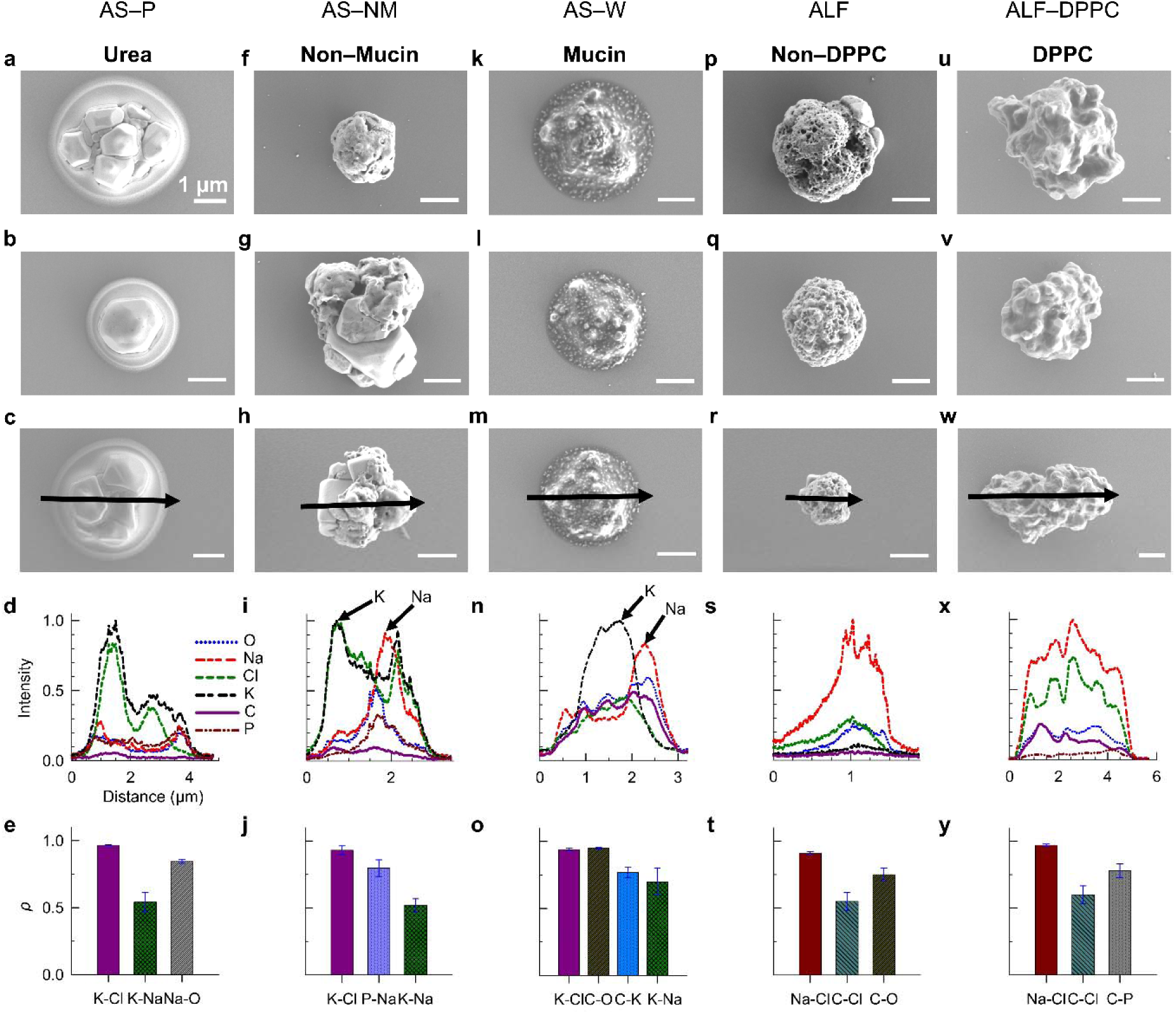
Organic composition (urea, mucin, and DPPC) modulates the morphology and organic-inorganic separation in aerosol particles derived from model respiratory fluids. Panels displa representative morphology, elemental distributions and Pearson correlation coefficients for artificial saliva from Pickering Lab (AS-P, 1700-0304) with urea as primary organic component, laboratory-prepared artificial saliva without mucin (AS-NM) and with mucin using Woo et al.’s formulation (AS-W), artificial lung fluid from Pickering Lab (ALF, 1700-0808) and ALF modified with dipalmitoylphosphatidylcholine (ALF-DPPC).

The presence of lipid such as DPPC leads to different aerosol morphologies for ALF. In absence of lipids, ALF produced dry, near-spherical particles with porous surfaces and, in many cases, small crystals are attached to core amorphous material (**Figure 3p-r & S21**). SEM and EDX analysis confirmed intermediate to high correlation coefficients among all element pairs, indicating that the components were well mixed overall (**Figure 3t**). When DPPC was introduced into ALF, the correlation coefficients remained comparable to those of lipid-free ALF, suggesting global mixing of elements. However, SEM images revealed more irregular, three-dimensional structures with smoother, more convex surfaces and pronounced dents. Multiple crystals appeared enclosed within other material (**Figure 3u-w, S22**). EDX line scans exhibited multiple peaks and minima of Na and Cl at corresponding positions, consistent with the presence of NaCl crystals and surface indentations. These results indicate that DPPC enhances surface restructuring and promotes crystal formation. Although the crystals are partially enclosed by organic material, localized phase separation becomes more pronounced compared to the more homogeneous and porous morphology observed in ALF-only particles.

### 3.4 Among all surrogate bioaerosol particles examined, AS-NM and EMEM complete media bioaerosol particles show size-dependent phase state

We further examined the size dependence of particle morphology by imaging particles collected across different stages of the Andersen impactor for both cell culture media and model respiratory fluids (**Figure 4**). Most virus-free bioaerosol particles, regardless of solution type, do not exhibit size-dependent morphologies, with similar structural features observed across stages (crystal, core-shell, phase separation, and well mixed). However, surrogate bioaerosol particles derived from EMEM complete medium and AS-NM show size-dependent phase state, where larger particles often appear semi-solid, showing spreading or splashing upon impaction, and smaller particles are dry and compact. This trend likely reflects differences in drying dynamics: under fixed RH and residence time, larger droplets can remain partially hydrated or semi-solid (Walker et al. 2021), while smaller droplets reach a dried-nuclei state more readily. Moreover, larger particles from semi-solid EMEM-complete medium frequently display core-shell structure, with multiple crystals encapsulated by an organic layer, while smaller desiccated particles appeared more homogeneous and well mixed.

**Figure 4.**
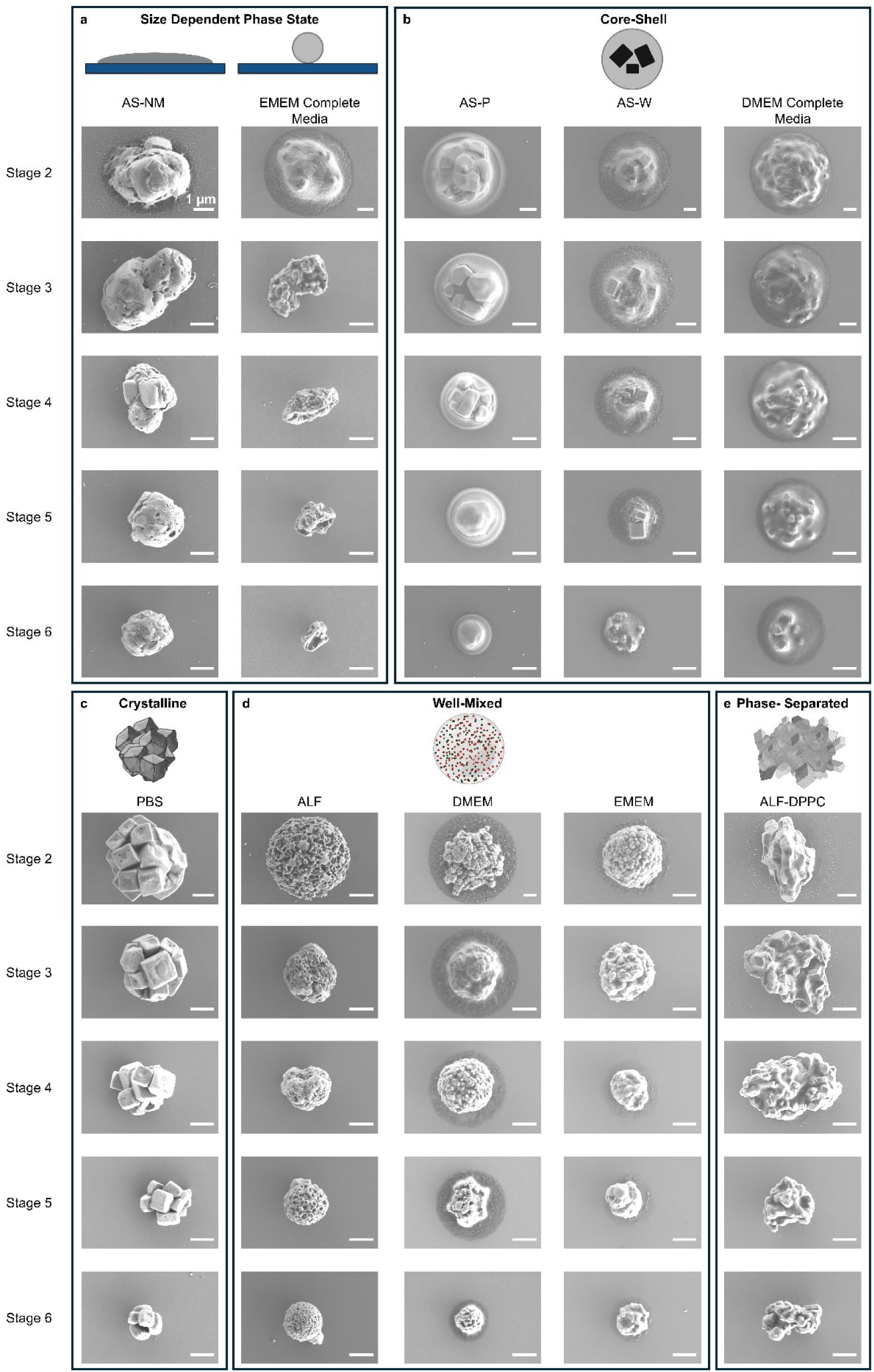
Among all surrogate bioaerosol particles examined, AS-NM and EMEM complete media show size-dependent phase state.

## 4. Discussions

### 4.1 Chemical distributions

The spatial distribution of chemical components within aerosol particles is an important factor because pathogen localization likely influences their survival during airborne suspension. As organic entities, pathogens may preferentially co-localize with organic-rich regions as suggested by observations in evaporating 1 µL droplets (Pan et al. 2025). Within aerosol particles, pathogen localization has not been experimentally examined and reported. Virus localization within aerosol particles has been hypothesized that viruses are localized with organics such as mucin (Alexander et al. 2022; Zuo et al. 2014; Woo et al. 2012) and BSA (Humphrey et al. 2023), thereby enhancing virus survival with increasing organic content. Nevertheless, if pathogen localization happens, chemical composition and evaporation kinetics will likely govern it. And reporting morphology can help uncover the links between virus survival and localization. The role of organic composition in determining chemical distributions is particularly significant. Proteins such as BSA and mucin promote compositional homogeneity and suppress the formation of large salt crystals, likely due to increased viscosity that restricts ionic diffusion. In contrast, organic solubility plays a dominant role in phase separation: glucose, being highly soluble, does not inhibit salt crystallization and produces core–shell morphologies in NaCl–glucose mixtures, whereas the poorly soluble lipid DPPC facilitates salt nucleation and promotes pronounced phase separation, leading to elongated or irregular particle shapes. These contrasting effects emphasize that both the molecular identity and physicochemical properties of organic components control aerosol morphology and chemical mixing. It is important to distinguish localized phase separation with the presence of small salt crystals when the chemical components appear globally well mixed. Chemical components exhibited intermediate to high Pearson correlation coefficients in particles generated from both cell culture media and model respiratory fluids, indicating globally well-mixed compositions. However, localized phase separation, with small salt crystals spatially separated from organics, was more pronounced in model respiratory fluids, while among cell culture media, only DMEM-complete medium exhibited similar localized separation.

Chemical distributions appeared largely independent of particle size within the 1–5 µm range investigated in this study. This observation contrasts with previous findings that sub-200 nm particles tend to exhibit homogeneous morphologies due to rapid evaporation (Altaf et al. 2016), whereas larger droplets undergo phase separation governed by the interplay between water loss and solute diffusion (Freedman 2020). The discrepancy likely arises from the greater chemical complexity and intermediate size range of the surrogate aerosols examined here.

### 4.2 Phase state: semi-solid

Aerosol phase state depends strongly on initial droplet size, chemical composition, relative humidity, and drying rate (Miles et al. 2025; Sapkota et al. 2025). For EMEM-complete medium, larger particles often remained semi-solid, whereas smaller ones were fully dried. Differences observed in AS-W particle morphologies compared with previous reports (Meng et al. 2025; Tian et al. 2024) are likely attributable to variations in both drying kinetics and mucin physicochemical properties. In the present study, particles experienced a residence time of approximately 10 s, which can limit water loss and favor semi-solid states, whereas longer residence time (∼50 s) shift the particles from semi-solid to solid morphologies consistent with reported morphologies (**Figure S23**). This difference in evaporation behavior might be due to the different types of mucins used as Type III porcine stomach mucin was used here while Type II mucin was employed in prior studies. For NaCl–glucose aerosols, previous studies have shown that salts such as NaCl depress the glass transition temperature and deliquescence RH of sugars (See et al. 2021), allowing glucose to remain semi-liquid at room temperature and low RH. Similarly, other organics may hinder water evaporation, maintaining semi-solid states in larger particles or under shorter residence times. Although a systematic analysis of drying kinetics was beyond the scope of this study, the results confirm that kinetic factors significantly influence virus-free bioaerosol phase behavior and morphology.

Although conventional SEM is not inherently suited for identifying semi-solid or viscous aerosol particles, combining impactor-based collection with projected-area equivalent diameter analysis enables indirect detection of semi-solid states. Semi-solid or viscous particles spread upon impaction due to high collection velocity within the impactor, forming circular boundaries visible in SEM images. In addition, particle cut-off sizes on different impactor stages are defined by the aerodynamic diameter (*d*□), which assumes spherical particles of unit density (1000 kg m□³). The expected particle diameter is typically smaller than the aerodynamic diameter for particles with a higher density than 1000 kg m□³. However, SEM measurements consistently showed projected-area equivalent diameters (*d_p_*) larger than the expected particle diameters for NaCl–glucose, DMEM-complete medium, AS-P, and AS-W particles across different ACI stages (**Figure S17**). This systematic deviation likely results from the spreading of semi-solid matrices upon impaction, which enlarges the projected area and inflates *d_p_* values.

## 5. Conclusions

This study demonstrates that coupling SEM imaging with EDX line scans and using an Andersen impactor for particle collection offer a powerful approach to investigate bioaerosol morphology and chemical distributions. This unique combination enables the visualization of three-dimensional structures and chemical spatial distributions not only for desiccated particles but also for semi-solid particles. It provides comprehensive insights into particle size (both aerodynamic and projected-area equivalent diameters), shape (spherical or irregular), internal chemical heterogeneity, and even phase variation between fully dried and semi-solid states. This method holds strong potential for the characterization of virus-laden bioaerosols in aerovirology research.

Our results show that aerosol morphology is highly dependent on the chemical composition of the aerosolization solutions, under the same drying conditions (∼30% RH and ∼10 s residence time). Distinct morphological categories were observed: (1) overall phase-separated particles with large salt crystals either exposed to or encapsulated by organics, (2) well-mixed particles with small, locally separated salt crystals, and (3) chemically homogeneous particles. In NaCl-organic systems, proteins such as BSA and mucin appeared to promote more uniform chemical distribution and suppress the formation of large crystals, likely due to increased solution viscosity. In contrast, low-solubility organics like DPPC, or highly soluble but non-viscous organics like glucose, tend to facilitate phase separation and the formation of large salt crystals.

Most particles reached a fully desiccated state under the study’s RH and residence time conditions (RH∼30% and residence time ∼10 s), though semi-solid particles were still observed, depending on the solution composition and particle size. For example, in EMEM-complete-media with high protein content, smaller particles were desiccated while larger particles remained semi-solid and showed spreading upon impaction. In contrast, DMEM-complete-media particles and NaCl-glucose exhibited semi-solid characteristics across all examined sizes, including NaCl-glucose nanoparticles (**Figure S24**), highlighting the strong influence of organic properties on phase behavior.

Finally, our findings underscore the importance of reporting virus-laden bioaerosol morphology and chemical composition in aerovirology studies. Morphology affects not only particle dynamics but also virus localization and survival. Caution should be taken when interpreting virus viability data across different studies, particularly when different aerosolization media are used. We recommend that future studies clearly report the chemical formulation of aerosolized solutions, particle size distributions, relative humidity, and drying times. Incorporating morphology and composition analyses will provide essential context for understanding airborne pathogen transmission and enhance the reproducibility and comparability of findings in the field.

## Acknowledgments

The authors acknowledge the Clean Room and Imaging Core Facility at the University of Texas at Dallas for providing access to SEM instrumentation.

## Disclosure Statement

No potential conflict of interest was reported by author(s).

## Funding

This work was supported by the National Institute of Allergy and Infectious Diseases of the National Institutes of Health under award numbers R21AI181258 and R21AI188518, as well as by startup funds from the University of Texas at Dallas.

## Notes

### Competing Interest Statement

The authors have declared no competing interest.

