## Supplementary material for "Organic Components Modulate the Morphology of Respirable Aerovirology-Relevant Aerosols": Supplimentary Information

**Table S1: Chemical Composition of EMEM.**

| **Chemical Compound** | **Concentration (g/L)** |
| --- | --- |
| CaCl_2_ (anhydrous) | 0.2 |
| KCl | 0.4 |
| MgSO_4_ (anhydrous) | 0.0977 |
| NaCl | 6.8 |
| NaH_2_PO_4_.H_2_O | 0.14 |
| NaHCO_3_ | 2.2 |
| L-Arginine.HCl | 0.1264 |
| L-Cystine .2HCl | 0.0312 |
| L-Glutamine | 0.292 |
| L-Histidine.HCl.H_2_O | 0.0419 |
| L-Isoleucine | 0.0525 |
| L-Leucine | 0.0525 |
| L-Lysine. HCl | 0.0725 |
| L-Methionine | 0.015 |
| L-Phenylalanine | 0.0325 |
| L-Threonine | 0.0476 |
| L-Tryptophan | 0.01 |
| L-Tyrosine.2Na.2H_2_O | 0.0519 |
| L-Valine | 0.0468 |
| D-Calcium pantothenate | 0.001 |
| Choline chloride | 0.001 |
| Folic acid | 0.001 |
| i-Inositol | 0.002 |
| Nicotinamide | 0.001 |
| Pyridoxine. HCl | 0.001 |
| Riboflavin | 0.0001 |
| Thiamine.HCl | 0.001 |
| D-Glucose | 1 |
| Phenol red.Na | 0.01 |
| **Total organic concentration** | 1.9 |
| **Total salt concentration** | 9.8 |
| **Organic-salt mass ratio (OSR)** | 0.19 |

**Table S2: Chemical Composition of DMEM.**

| **Chemical Compound** | **Concentration (g/L)** |
| --- | --- |
| CaCl_2_ (anhydrous) | 0.2 |
| Fe(NO_3_)_3_.9H_2_O | 0.0001 |
| KCl | 0.400 |
| MgSO₄ (anhydrous) | 0.0977 |
| NaCl | 6.400 |
| NaH₂PO₄.H₂O | 0.125 |
| NaHCO₃ | 3.700 |
| L-Arginine.HCl | 0.084 |
| L-Cystine.2HCl | 0.06257 |
| L-Glutamine | 0.584 |
| Glycine | 0.030 |
| L-Histidine.HCl.H₂O | 0.042 |
| L-Isoleucine | 0.1048 |
| L-Leucine | 0.1048 |
| L-Lysine.HCl | 0.1462 |
| L-Methionine | 0.030 |
| L-Phenylalanine | 0.066 |
| L-Serine | 0.042 |
| L-Threonine | 0.0952 |
| L-Tryptophan | 0.016 |
| L-Tyrosine.2Na.2H₂O | 0.10379 |
| L-Valine | 0.094 |
| D-Calcium Pantothenate | 0.004 |
| Choline Chloride | 0.004 |
| Folic Acid | 0.004 |
| i-Inositol | 0.0072 |
| Nicotinamide | 0.004 |
| Pyridoxine.HCl | 0.004 |
| Riboflavin | 0.0004 |
| Thiamine.HCl | 0.004 |
| D-Glucose | 4.500 |
| Phenol Red.Na | 0.015 |
| **Total organic concentration** | 6.14 |
| **Total salt concentration** | 10.94 |
| **Organic-salt mass ratio (OSR)** | 0.56 |

**Table S3: Chemical Composition of DMEM Complete Media**

| **Chemical Compound** | **Concentration (g/L)** |
| --- | --- |
| CaCl_2_ (anhydrous) | 0.17 |
| Fe(NO_3_)_3_.9H_2_O | 0.00008 |
| KCl | 0.33 |
| MgSO₄ (anhydrous) | 0.08 |
| NaCl | 5.33 |
| NaH₂PO₄.H₂O | 0.1042 |
| NaHCO₃ | 3.083 |
| L-Arginine.HCl | 0.07 |
| L-Cystine.2HCl | 0.05 |
| L-Glutamine | 0.487 |
| Glycine | 0.025 |
| L-Histidine.HCl.H₂O | 0.035 |
| L-Isoleucine | 0.0873 |
| L-Leucine | 0.0873 |
| L-Lysine.HCl | 0.1218 |
| L-Methionine | 0.025 |
| L-Phenylalanine | 0.055 |
| L-Serine | 0.035 |
| L-Threonine | 0.079 |
| L-Tryptophan | 0.0133 |
| L-Tyrosine.2Na.2H₂O | 0.087 |
| L-Valine | 0.078 |
| D-Calcium Pantothenate | 0.0033 |
| Choline Chloride | 0.0033 |
| Folic Acid | 0.0033 |
| i-Inositol | 0.006 |
| Nicotinamide | 0.0033 |
| Pyridoxine.HCl | 0.0033 |
| Riboflavin | 0.0003 |
| Thiamine.HCl | 0.0033 |
| Glucose | 3.81 |
| Phenol Red.Na | 0.0125 |
| FBS Protein* | 3.5 |
| Urea* | 0.012 |
| Penicillin* | 0.001 |
| **Total organic concentration** | 8.7 |
| **Total salt concentration** | 9.1 |
| **Organic-salt mass ratio (OSR)** | 0.96 |

***Estimation from FBS solution (Fisherbrand Research Grade Fetal Bovine Serum, Canadian Sourced, Cat. No: FB12999102)**

**Table S4: Chemical Composition of EMEM Complete Media**

| **Chemical Compound** | **Concentration (g/L)** |
| --- | --- |
| CaCl_2_ (anhydrous) | 0.17 |
| KCl | 0.33 |
| MgSO₄ (anhydrous) | 0.08 |
| NaCl | 5.66 |
| NaH₂PO₄.H₂O | 0.117 |
| NaHCO₃ | 1.83 |
| L-Arginine.HCl | 0.105 |
| L-Cystine.2HCl | 0.026 |
| L-Glutamine | 0.487 |
| L-Histidine.HCl.H₂O | 0.035 |
| L-Isoleucine | 0.0435 |
| L-Leucine | 0.060 |
| L-Lysine.HCl | 0.1218 |
| L-Methionine | 0.0125 |
| L-Phenylalanine | 0.027 |
| L-Serine | 0.035 |
| L-Threonine | 0.0397 |
| L-Tryptophan | 0.0083 |
| L-Tyrosine.2Na.2H₂O | 0.04325 |
| L-Valine | 0.039 |
| D-Calcium Pantothenate | 0.00083 |
| Choline Chloride | 0.00083 |
| Folic Acid | 0.00083 |
| i-Inositol | 0.0017 |
| Nicotinamide | 0.00083 |
| Pyridoxine.HCl | 0.00083 |
| Riboflavin | 0.00008 |
| Thiamine.HCl | 0.00083 |
| Glucose | 0.897 |
| Phenol Red.Na | 0.0083 |
| FBS Protein* | 3.5 |
| Urea* | 0.012 |
| Penicillin* | 0.066 |
| **Total organic concentration** | 5.21 |
| **Total salt concentration** | 8.20 |
| **Organic-salt mass ratio (OSR)** | 0.64 |

***Estimation from FBS solution (Fisherbrand Research Grade Fetal Bovine Serum, Canadian Sourced, Cat. No: FB12999102)**

**Table S5: Chemical composition of phosphate buffered saline.**

| **Chemical Compound** | **Concentration (g/L)** |
| --- | --- |
| KH_2_PO_4_ | 0.144 |
| NaCl | 9 |
| Na_2_HPO_4_ (anhydrous) | 0.795 |
| **Total organic concentration** | 0 |
| **Total salt concentration** | 9.94 |
| **Organic-salt mass ratio (OSR)** | 0 |

**Table S6: Chemical Composition of Artificial Saliva Pickering Lab CA USA (AS–P, 1700-0304)**

| **Chemical Compound** | **Concentration (g/L)** |
| --- | --- |
| NaCl | 0.126 |
| KH_2_PO_4_ | 0.655 |
| KCl | 0.964 |
| KSCN | 0.189 |
| (NH_2_)_2_CO (Urea) | 0.20 |
| **Total organic concentration** | 0.2 |
| **Total salt concentration** | 1.93 |
| **Organic-salt mass ratio (OSR)** | 0.1 |

**Table S7: Chemical Composition of Artificial Saliva Non-Mucin (AS–NM)**

| **Chemical Compound** | **Concentration (g/L)** |
| --- | --- |
| MgCl_2_.7H_2_O | 0.04 |
| CaCl_2_.H_2_O | 0.13 |
| NaHCO_3_ | 0.42 |
| KH_2_PO_4_ | 0.2096 |
| K_2_HPO_4_ | 0.4285 |
| KCl | 1.04 |
| NH_4_Cl | 0.11 |
| KSCN | 0.19 |
| (NH_2_)_2_CO | 0.12 |
| NaCl | 0.88 |
| DMEM | 0.001 |
| Mucin | - |
| **Total organic concentration** | 0.12 |
| **Total salt concentration** | 3.45 |
| **Organic-salt mass ratio (OSR)** | 0.04 |

**Table S8: Chemical Composition of Artificial Saliva Woo’s Formulation (AS–W)**

| **Chemical Compound** | **Concentration (g/L)** |
| --- | --- |
| MgCl_2_.7H_2_O | 0.04 |
| CaCl_2_.H_2_O | 0.13 |
| NaHCO_3_ | 0.42 |
| KH_2_PO_4_ | 0.2096 |
| K_2_HPO_4_ | 0.4285 |
| KCl | 1.04 |
| NH_4_Cl | 0.11 |
| KSCN | 0.19 |
| (NH_2_)_2_CO | 0.12 |
| NaCl | 0.88 |
| DMEM | 0.001 |
| Porcine Stomach Mucin | 3 |
| **Total organic concentration** | 3.12 |
| **Total salt concentration** | 3.45 |
| **Organic-salt mass ratio (OSR)** | 0.91 |

**Table S9: Chemical Composition of Artificial Lung Fluid Pickering Lab CA USA (1700-0808) in solution.**

| **Chemical Compound** | **Concentration (g/L)** |
| --- | --- |
| MgCl_2_.6H_2_O | 0.203 |
| NaCl | 6.019 |
| KCl | 0.298 |
| Na_2_HPO_4_ | 0.126 |
| Na_2_SO_4_.10H_2_O | 0.681 |
| CaCl_2_.2H_2_O | 0.368 |
| NaHCO_3_ | 2.604 |
| C_6_H_5_Na_3_O_7_.2H_2_O | 0.097 |
| C_2_H_3_O_2_Na.3H_2_O | 0.952 |
| Pro-Clean (Anti-Bacterial Solution) | 0.3 ml/L |
| **Total organic concentration** | 0 |
| **Total salt concentration** | 11.35 |
| **Organic-salt mass ratio (OSR)** | 0 |

**Table S10: Chemical Composition of modified Artificial Lung Fluid with DPCC**

| **Chemical Compound** | **Concentration (g/L)** |
| --- | --- |
| MgCl_2_.6H_2_O | 0.05 |
| NaCl | 1.5 |
| KCl | 0.075 |
| Na_2_HPO_4_ | 0.0315 |
| Na_2_SO_4_.10H_2_O | 0.170 |
| CaCl_2_.2H_2_O | 0.092 |
| NaHCO_3_ | 0.651 |
| C_6_H_5_Na_3_O_7_.2H_2_O | 0.024 |
| C_2_H_3_O_2_Na.3H_2_O | 0.238 |
| Pro-Clean (Anti-Bacterial Solution) | 0.3 ml/L |
| DPPC | 1.2 |
| **Total organic concentration** | 1.2 |
| **Total salt concentration** | 2.84 |
| **Organic-salt mass ratio (OSR)** | 0.42 |

**Table S11: Aerodynamic Cut-Off diameter of different stages of Andersen Impactor at 28.3 L/min flow rate.**

| **Stage** | **Aerodynamic Cut-Off Diameter at 28.3 LPM (** $\boldsymbol{d}_{\boldsymbol{ae}}$ **)(µm)** |
| --- | --- |
| 2 | 4.7-5.8 |
| 3 | 3.3-4.7 |
| 4 | 2.1-3.3 |
| 5 | 1.1-2.1 |
| 6 | 0.7-1.1 |

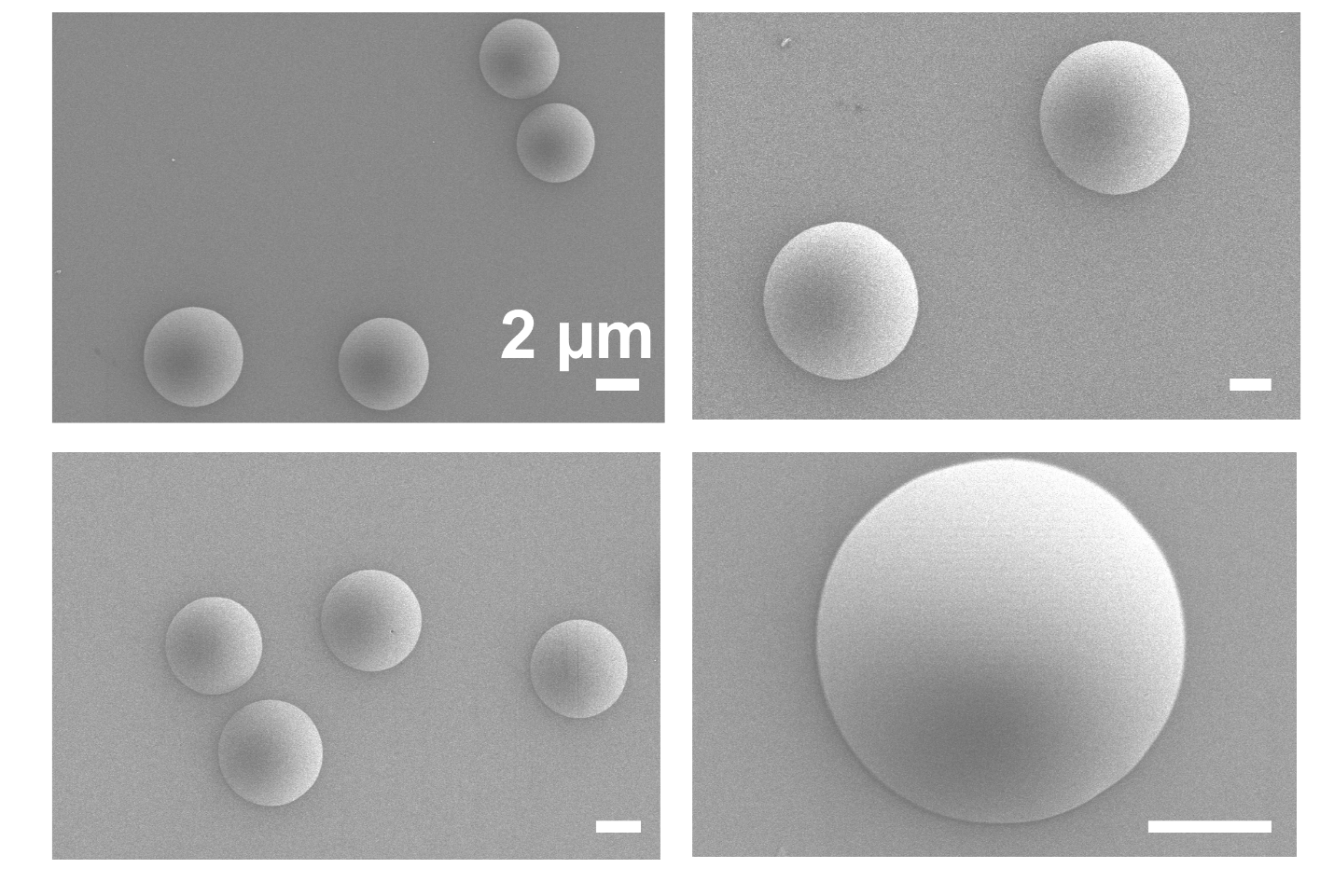

**Figure S1. SEM image s of glucose (20 g/L) aerosol particles.**

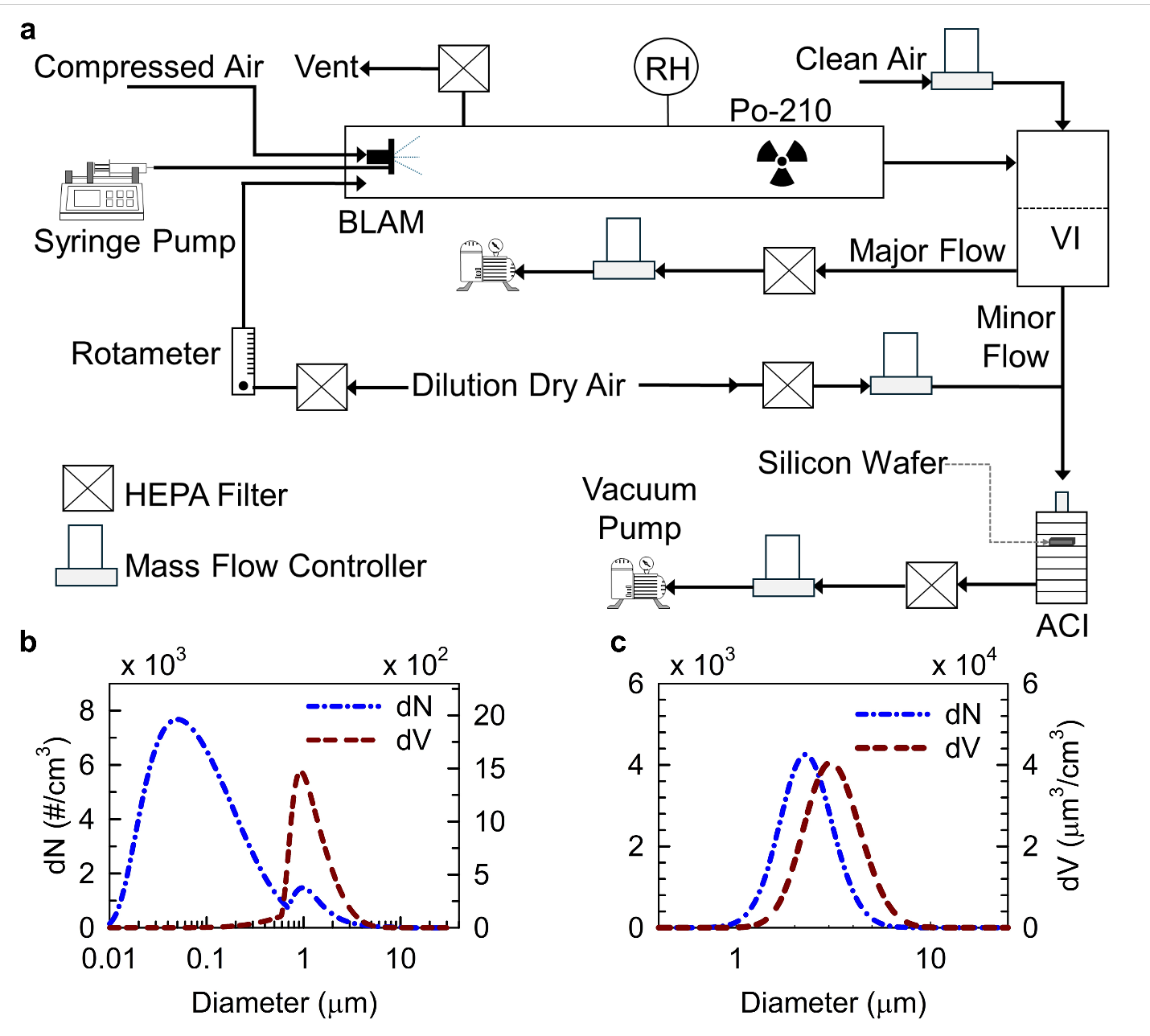

**Figure S2. Experimental setup and size distribution of bioaerosol particles.** (a) Schematic diagram illustrating the bioaerosol generation and collection process. (b) and (c) Particle size distributions (PSD) of DMEM aerosols based on both number concentration (dN, #/cm^3^, left) and volume concentration (dV, µm³/ cm^3^, right), measured at the inlet (b) and exit (c) of the virtual impactor (VI). Abbreviations: RH, relative humidity; SMPS, Scanning Mobility Particle Sizer; APS, Aerodynamic Particle Spectrometer.

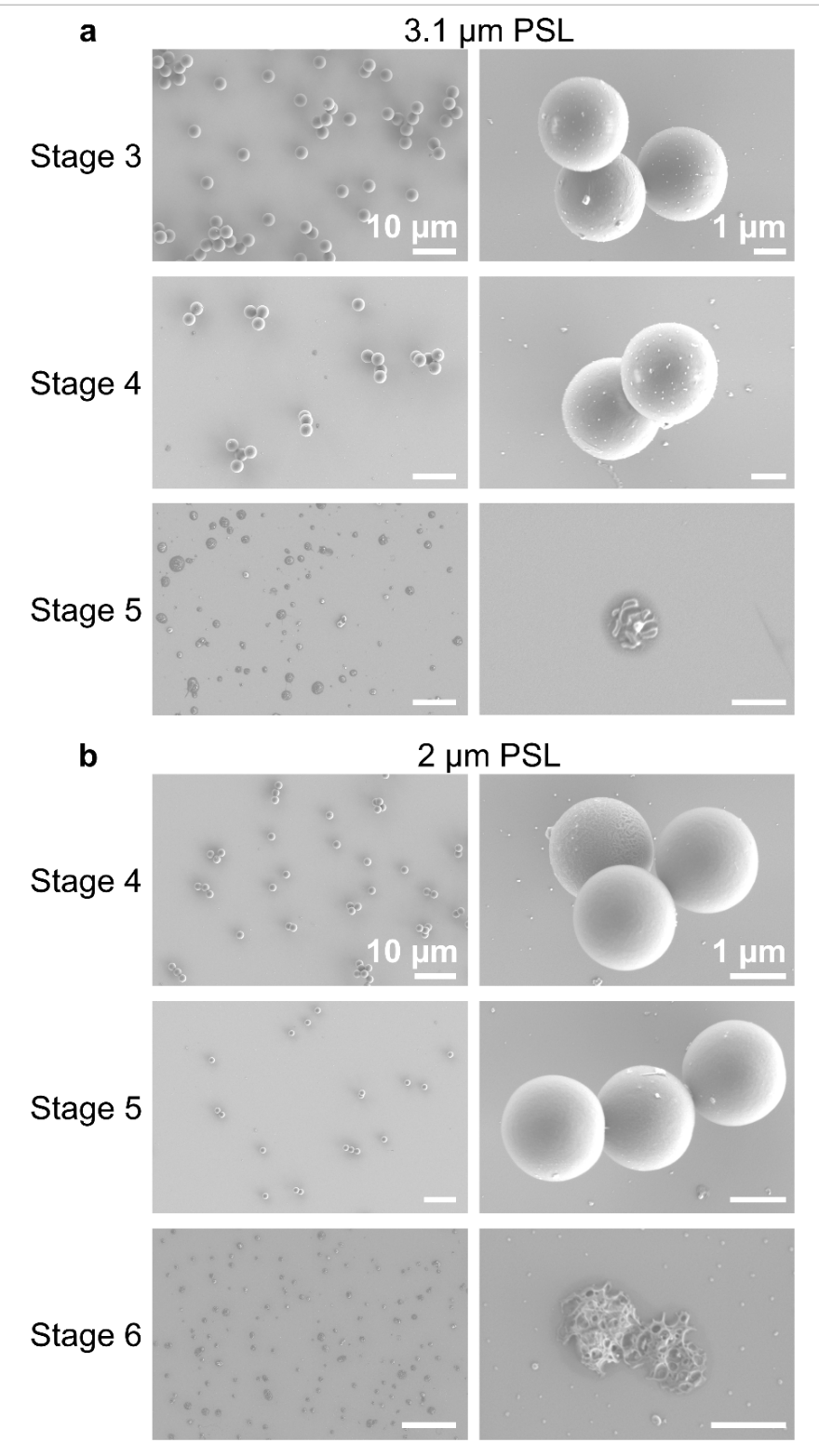

**Figure S3. SEM images of polystyrene latex sphere (PSL) particles collected at various stages of the Andersen Cascade Impactor operated at a standardized flow rate of 28.3 L/min.** Note: The flow rate was held constant for all subsequent figures unless specified otherwise.

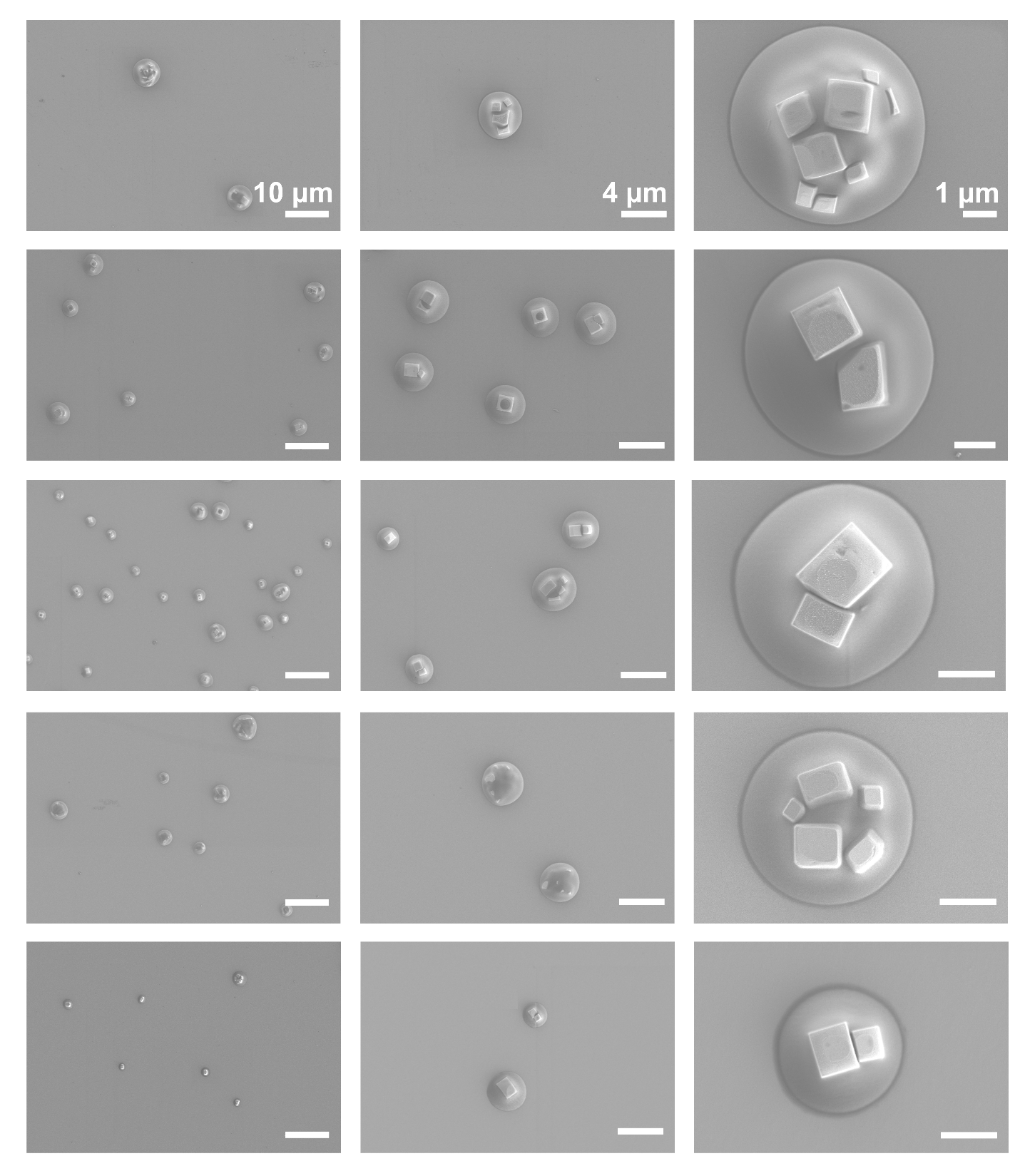

**Figure S4. Two-component NaCl–glucose aerosols show core-shell morphology.** Particles are collected at different stages of the Andersen cascade impactor. Mass concentration is 6 g/L for both NaCl and glucose.

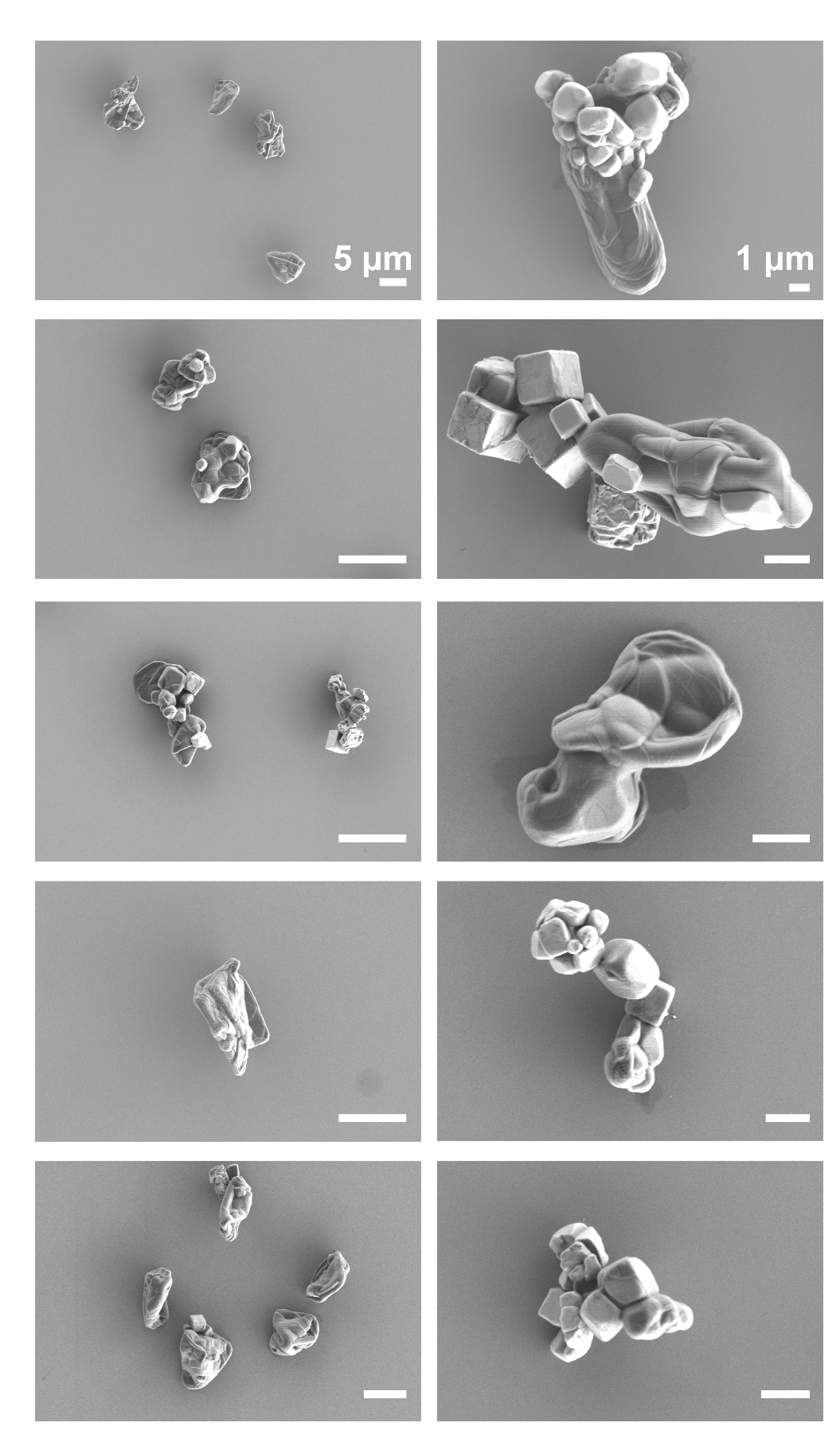

**Figure S5. SEM images for NaCl****–DPPC aerosols.** The concentrations for both NaCl and DPPC are 6 g/L.

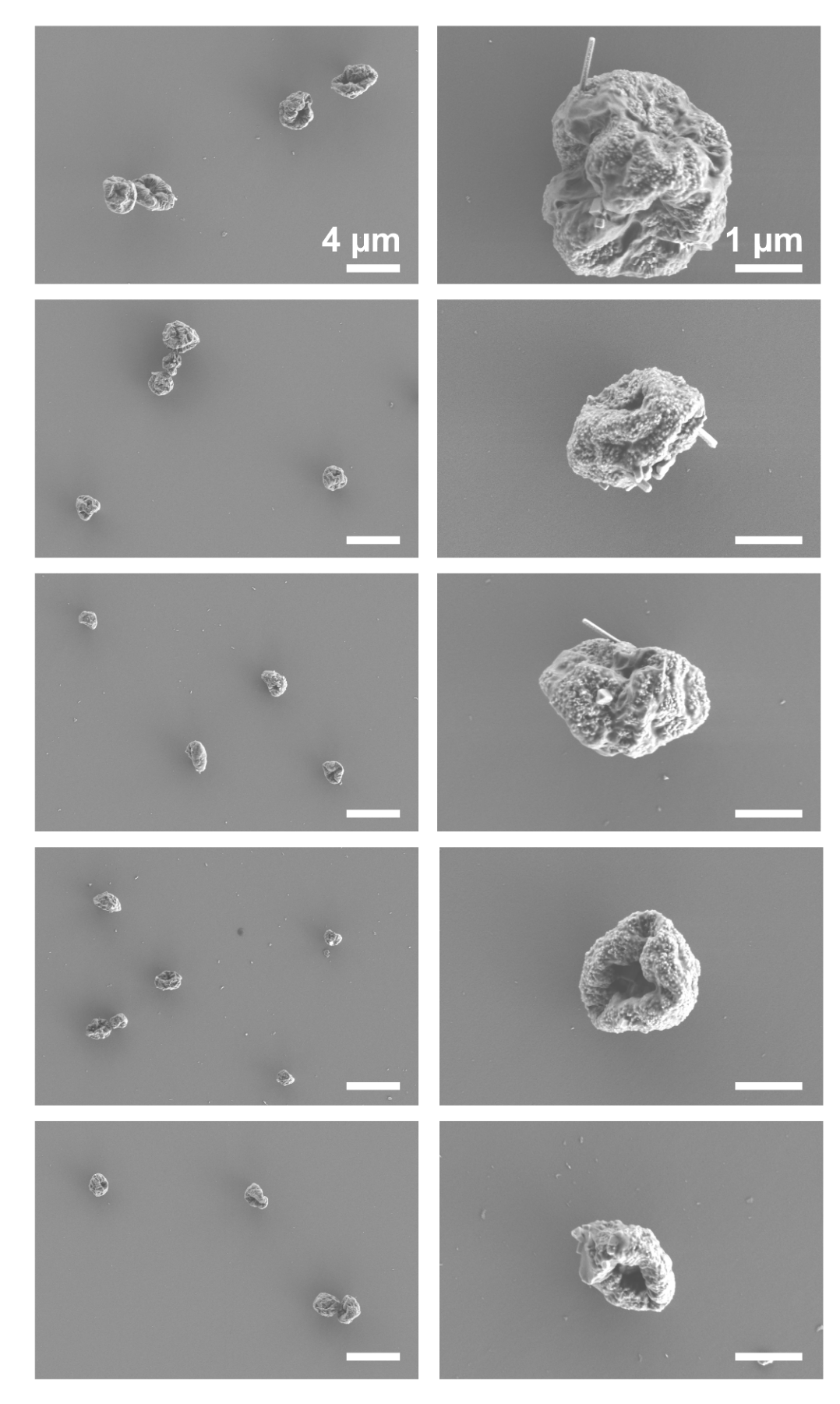

**Figure S6. SEM images for NaCl–Mucin aerosols.** The concentrations for both NaCl and mucin are 6 g/L.

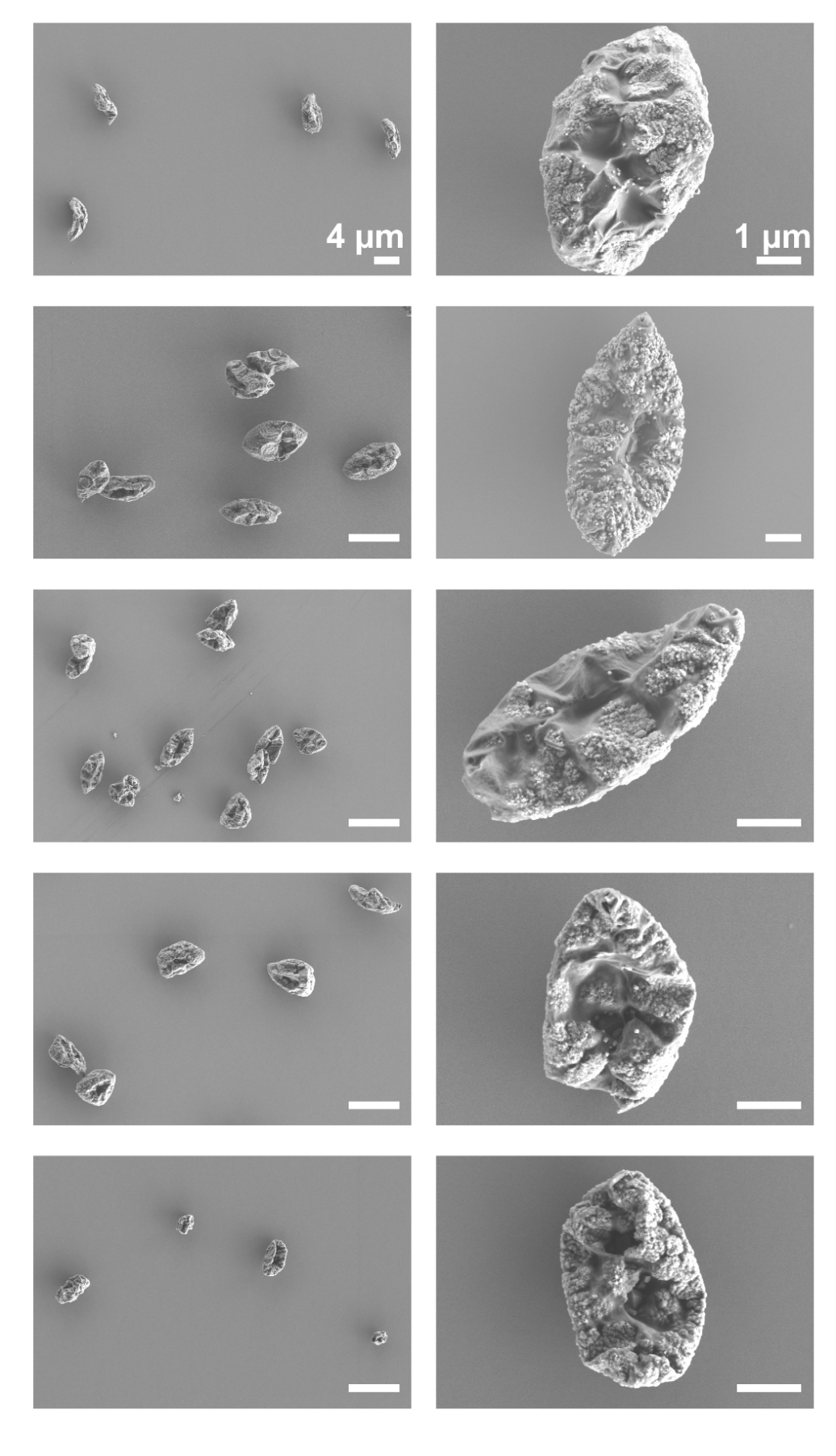

**Figure S7. SEM images for NaCl–BSA aerosols.** The concentrations for both NaCl and BSA are 6 g/L.

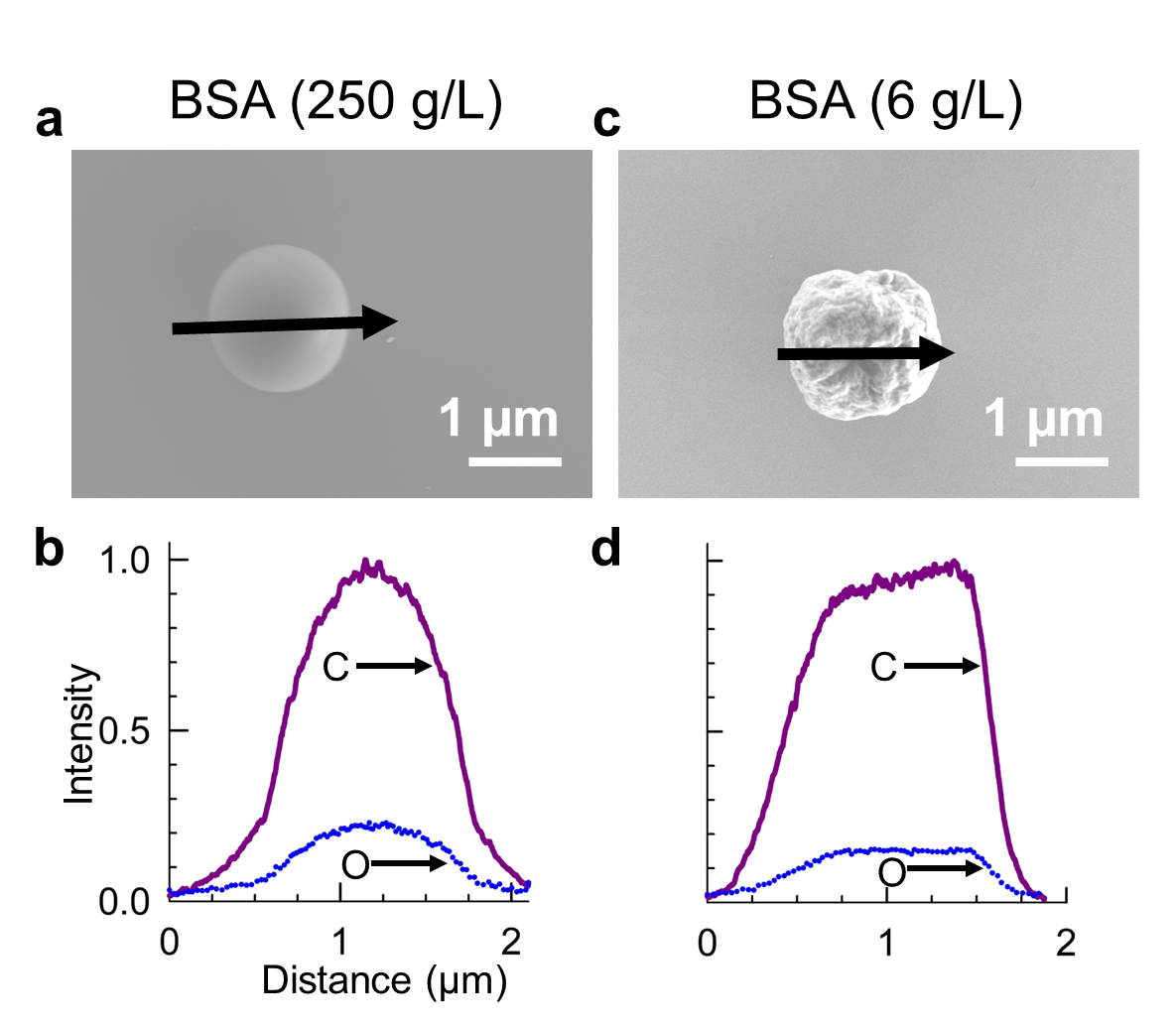
**Figure S8. SEM image and corresponding EDX line scan of BSA aerosol particles with concentration of (a-b) 250 g/L and (c-d) 6 g/L.**

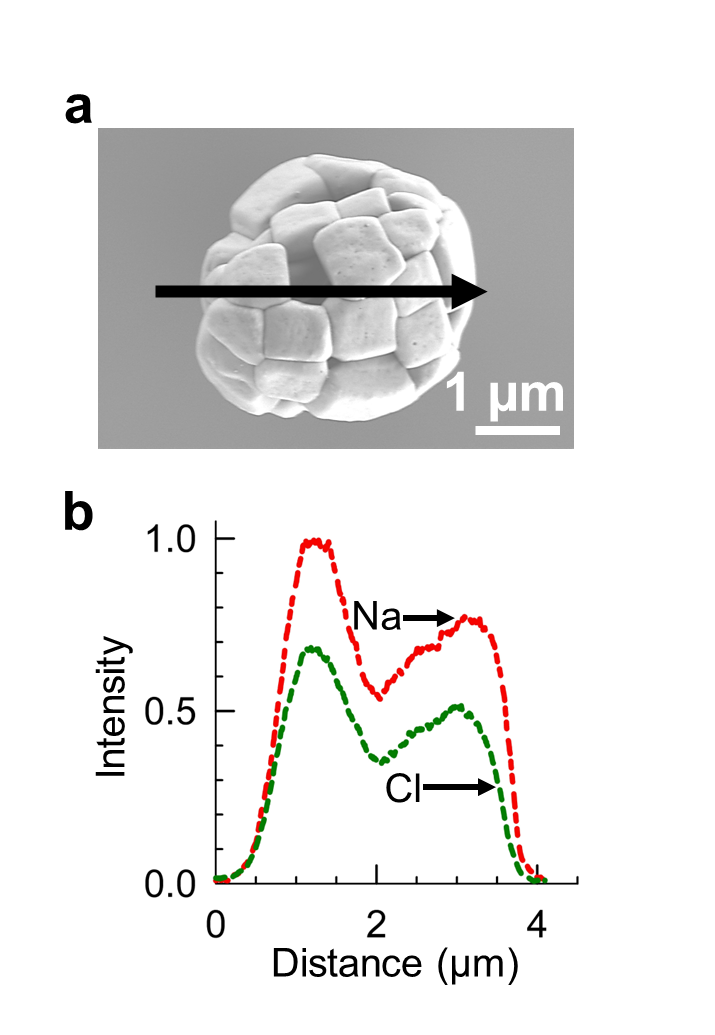

**Figure S9. SEM image (a) and EDX line scan (b) of NaCl only aerosol particle.**

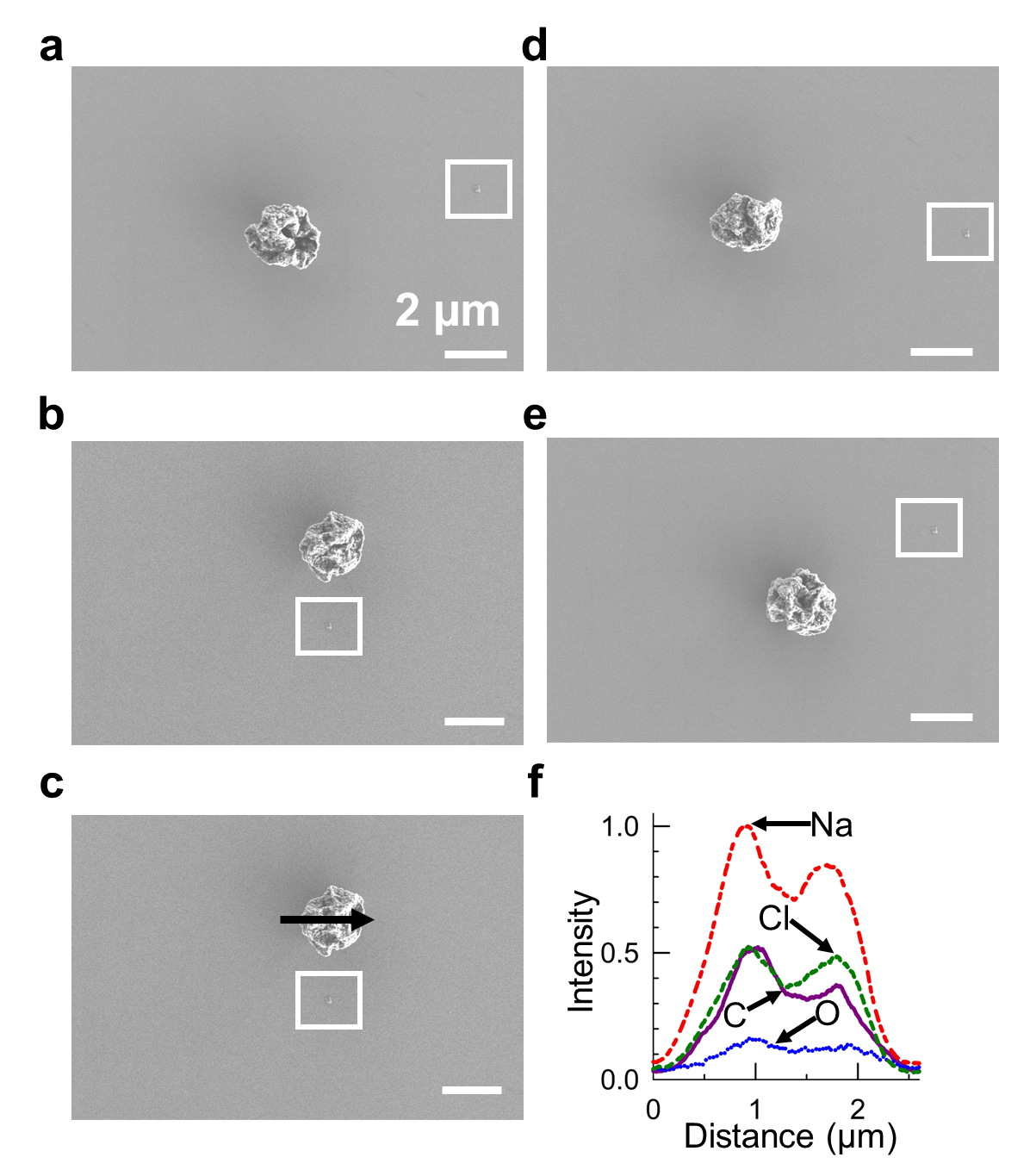

**Figure S10. SEM images and EDX line scan of a NaCl–BSA particle that flipped during SEM imaging.** The white rectangular box marks the reference position used to track particle orientation.

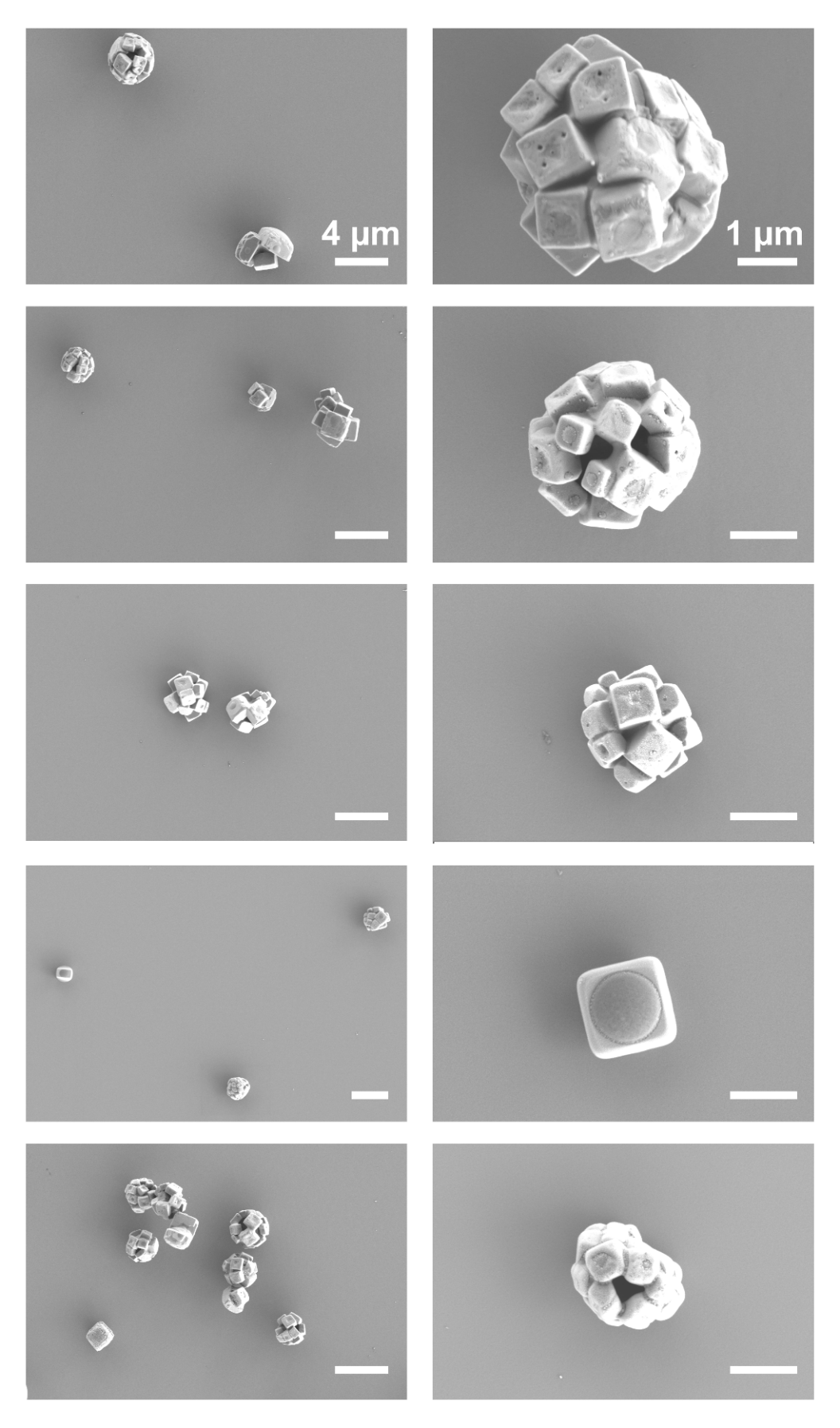

**Figure S11. SEM images for PBS aerosols.**

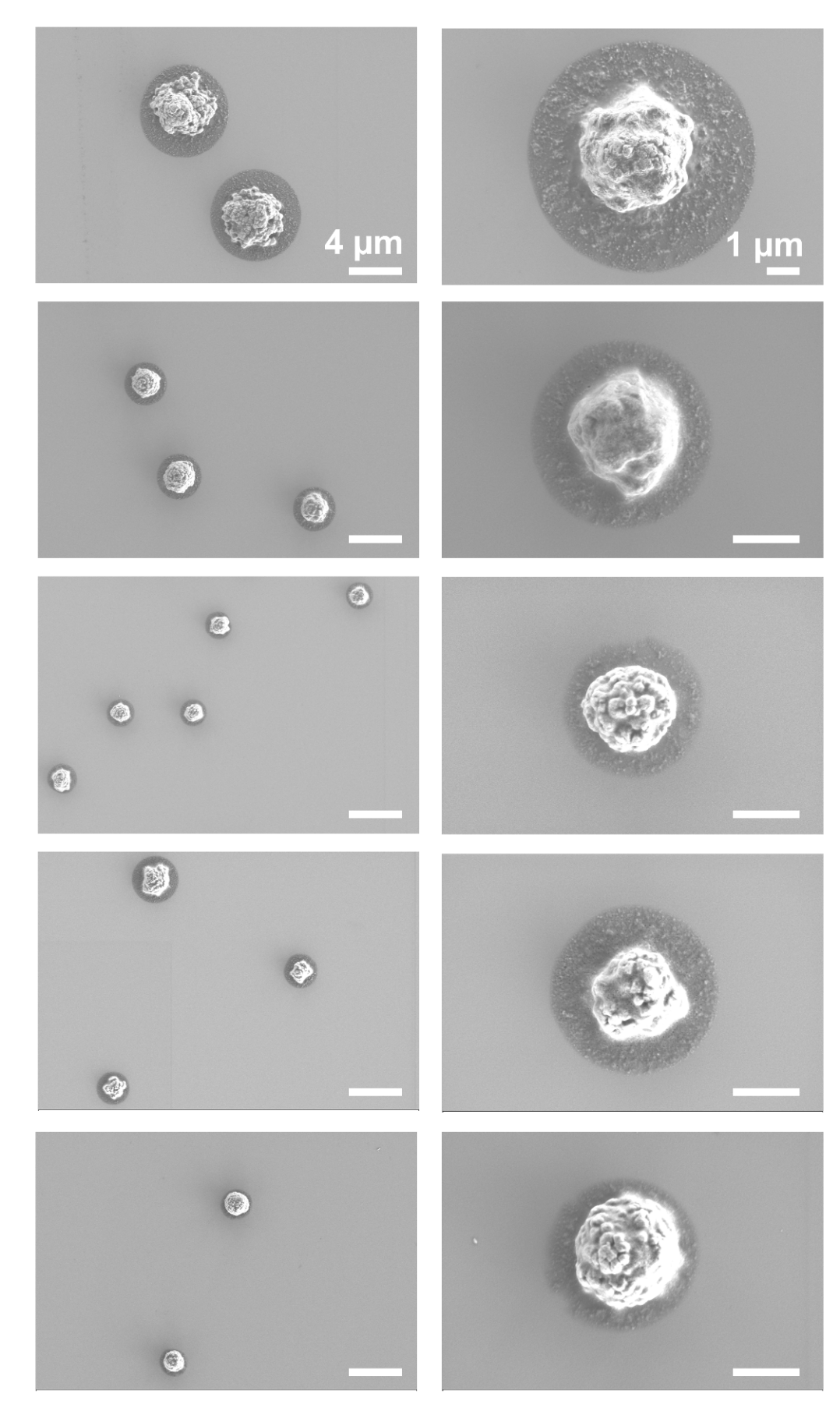

**Figure S12. SEM images for DMEM aerosols.**

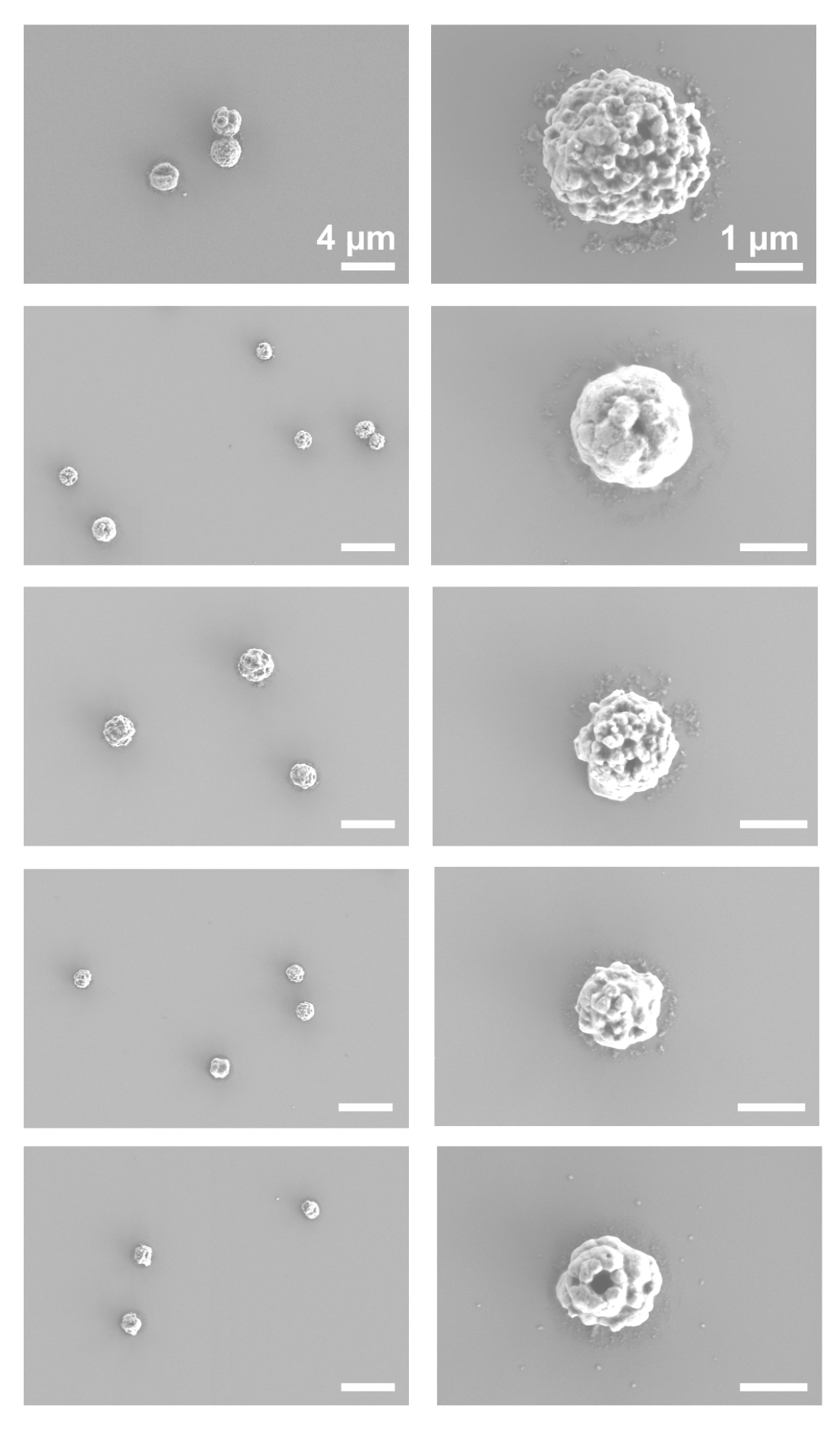

**Figure S13.SEM images for EMEM aerosols**

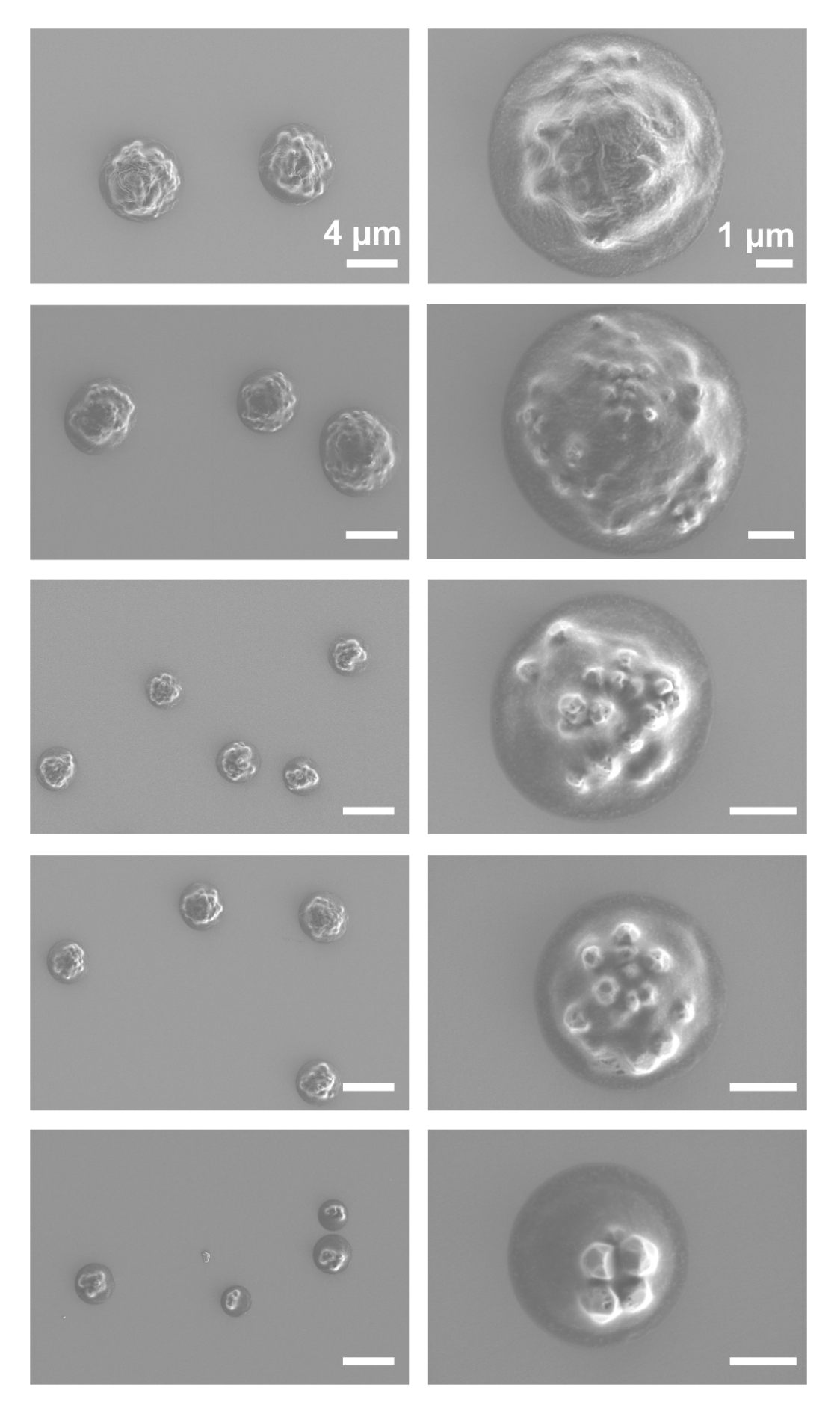

**Figure S14. SEM images for DMEM complete media aerosols.**

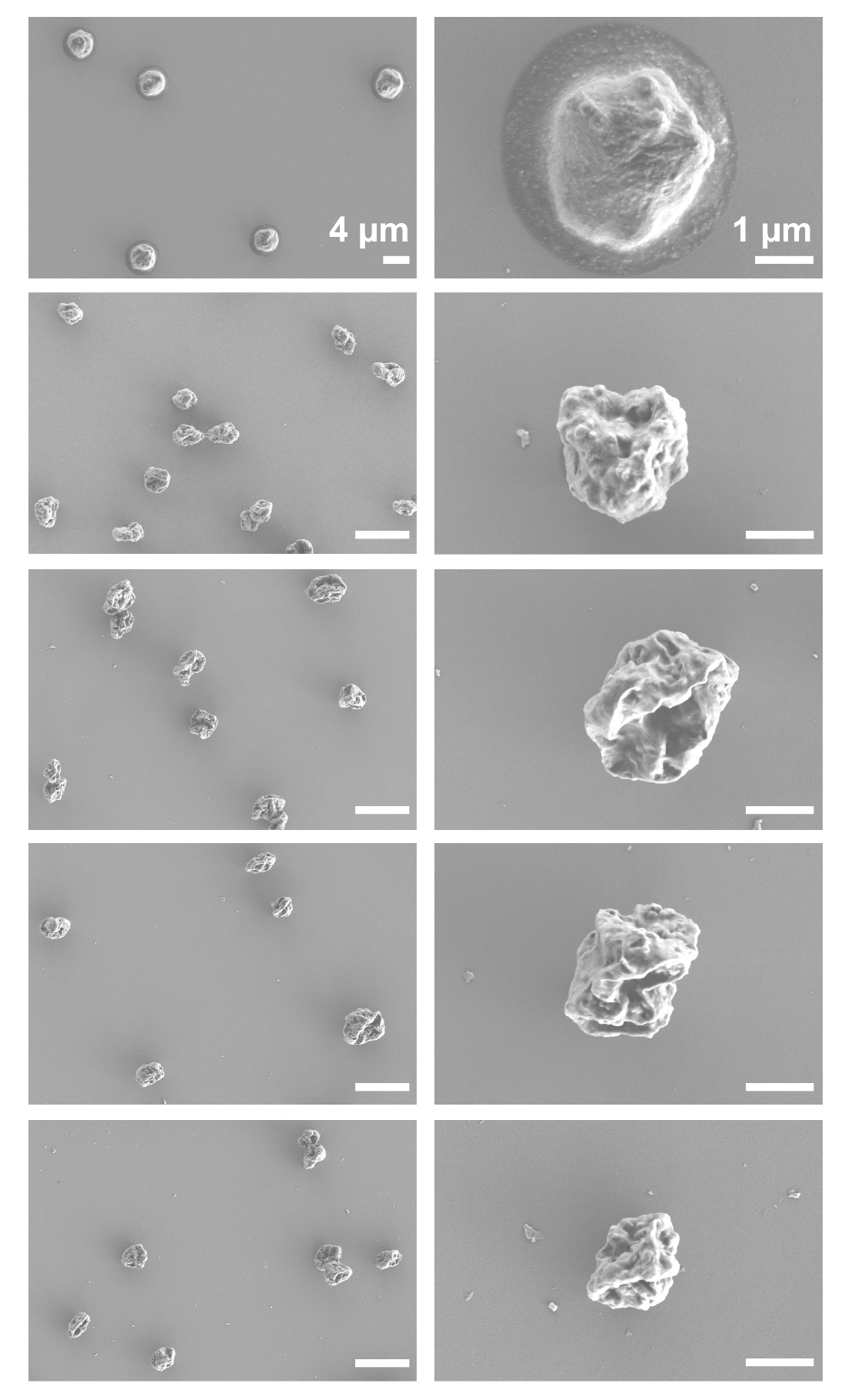

**Figure S15. SEM images for EMEM complete media aerosols**

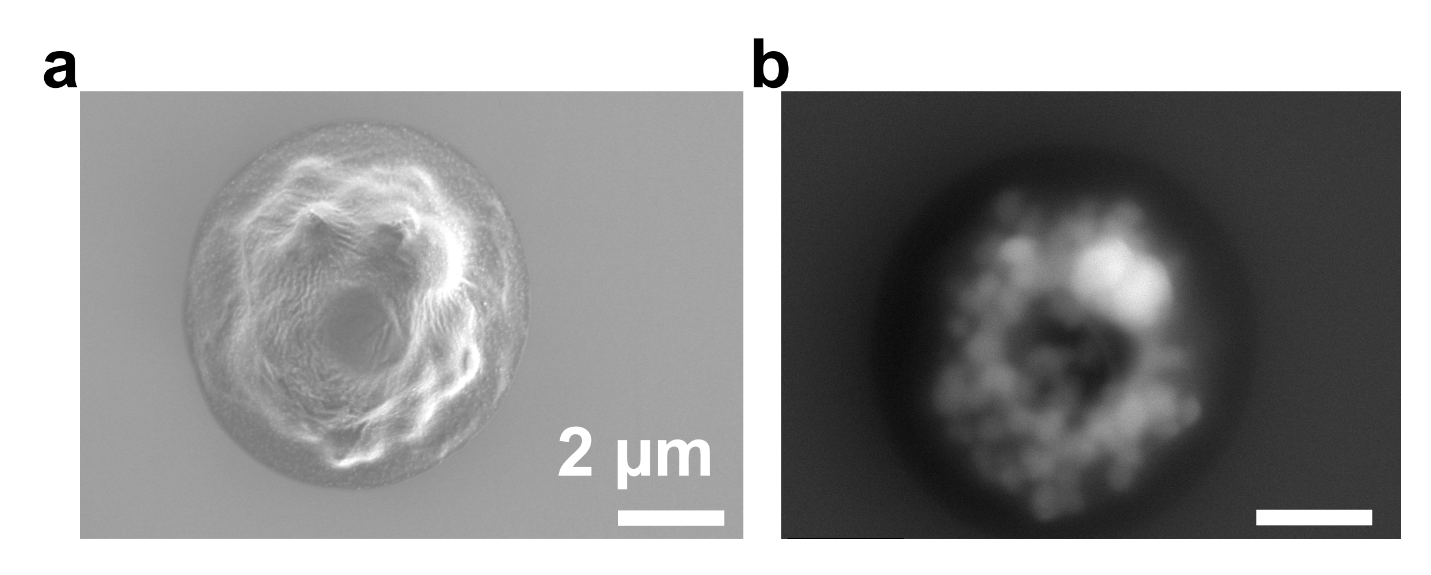
 **Figure S16. SEM images of DMEM Complete Media aerosol particle acquired under (a) low and (b) high acceleration voltages.** The low-voltage image (a) reveals surface morphology and features of the serum layer, while the high-voltage image (b) confirms the presence of salt crystals previously embedded within the serum matrix.

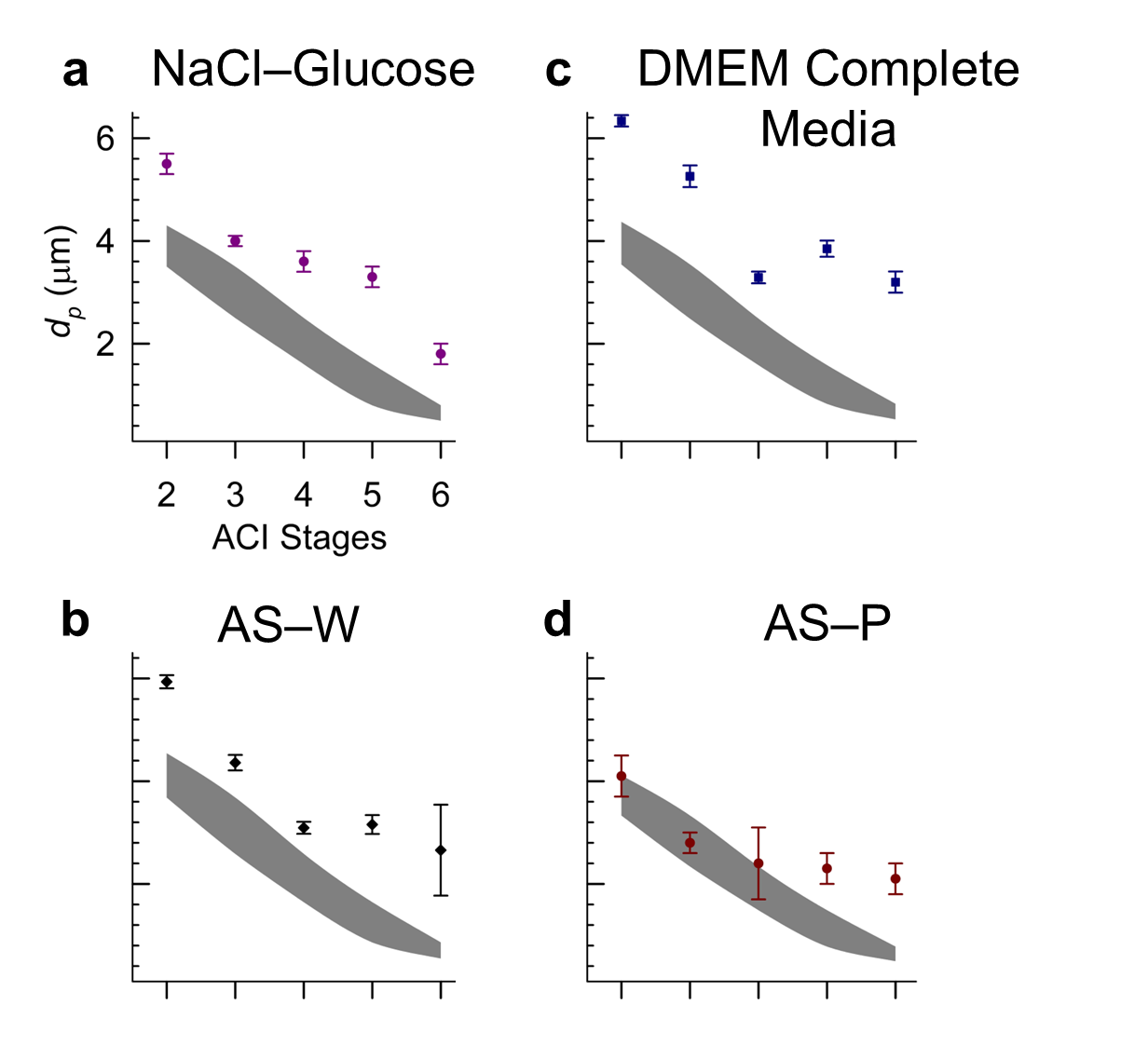

**Figure S17: Projected-area equivalent diameters (symbols) are consistently larger than expected particle diameter (grey region) across different ACI stages.** Projected area equivalent diameters are measured using ImageJ with SEM images while the expected particle diameters $d_{p}=d_{ae}\sqrt{\rho_{o}/\rho_{p}}$ are calculated from the aerodynamic diameter ($d_{ae}$) assuming reference water density *ρ_o_* = 1 g/cc and estimated particle densities *ρ_p_* of : (a) 1.81 g/cc, (b) 1.63 g/cc, (c) 1.76 g/cc, and (d) 1.99 g/cc.

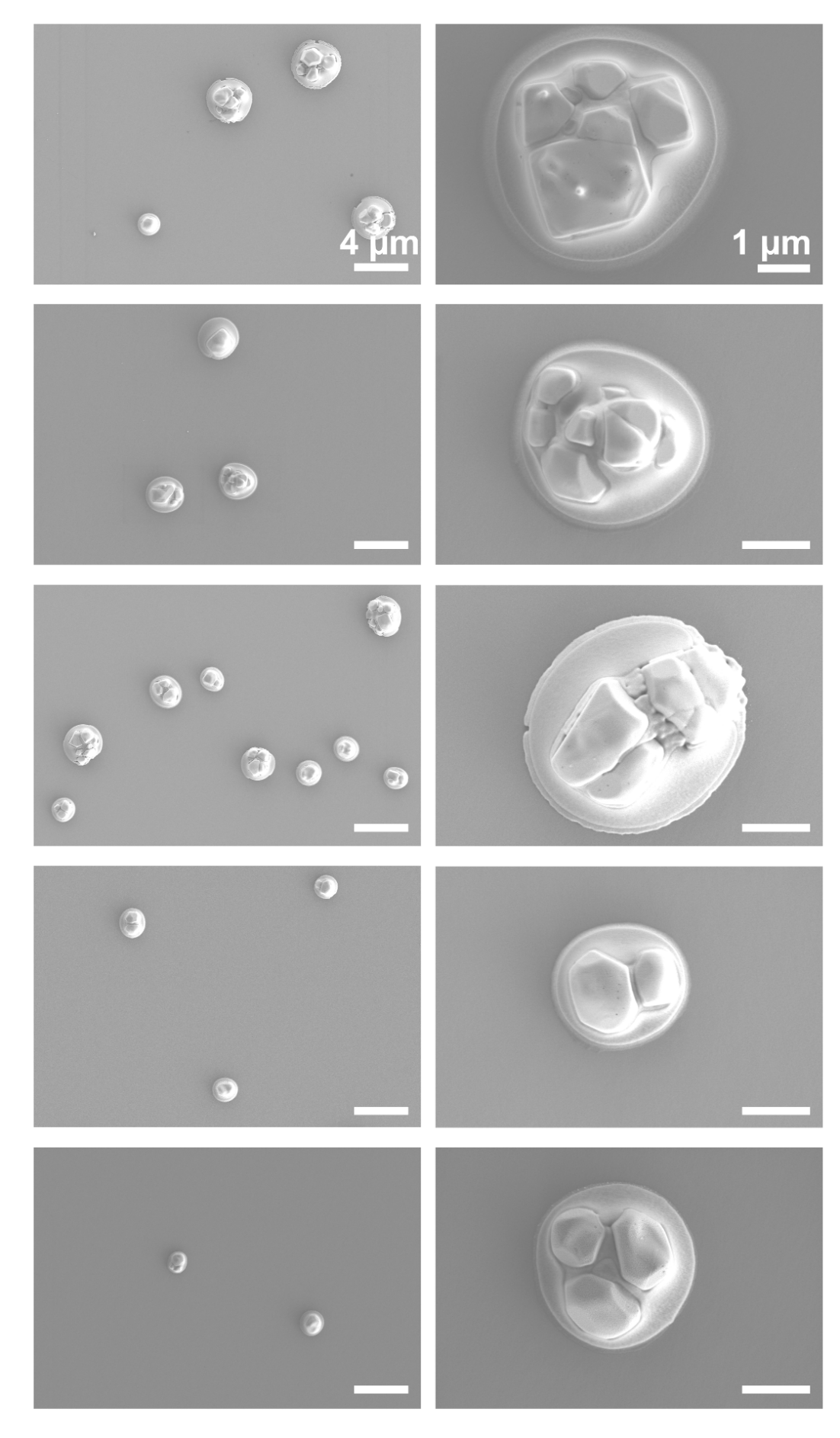

**Figure S18. SEM images for AS–P aerosols**

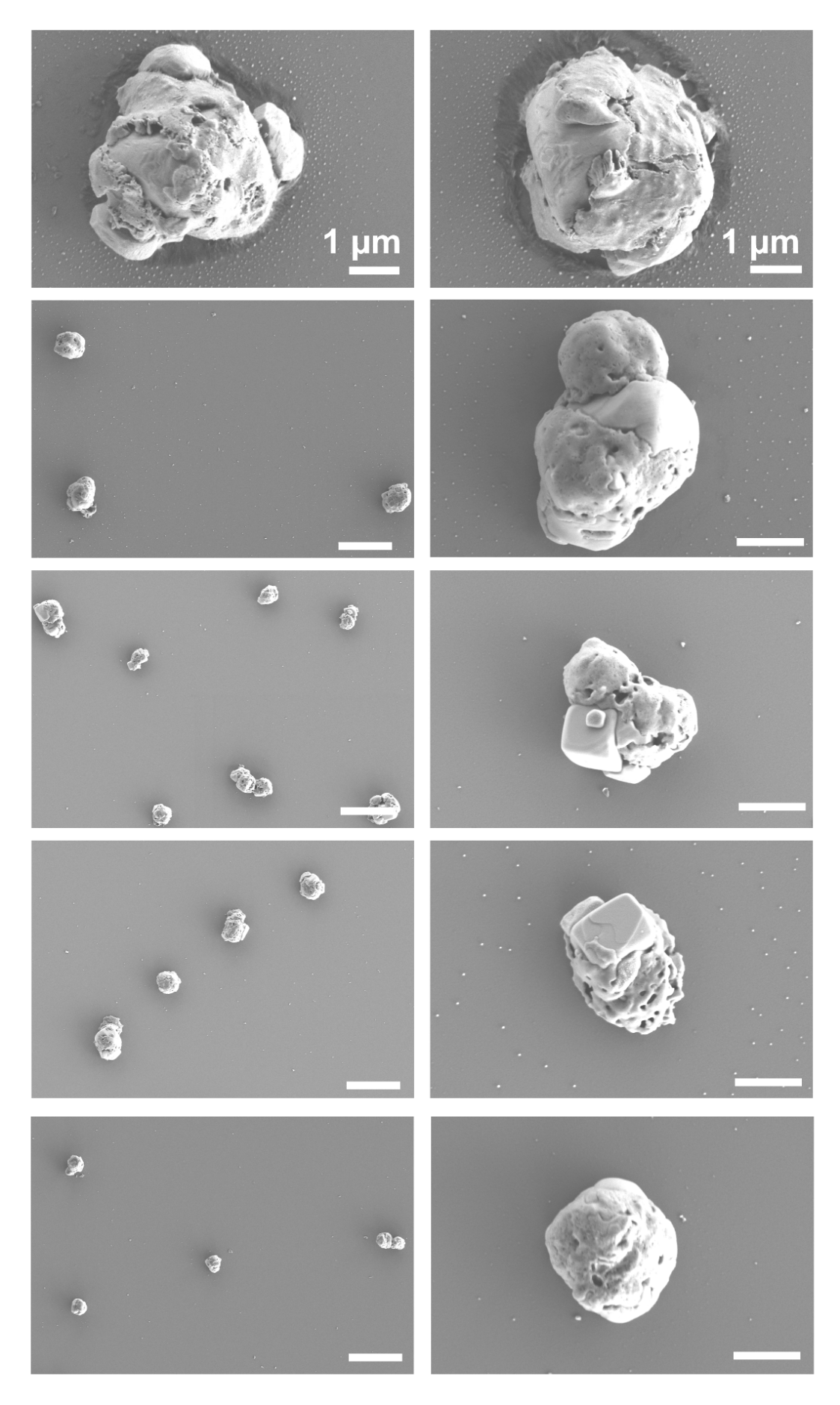

**Figure S19. SEM images for AS–NM aerosols**

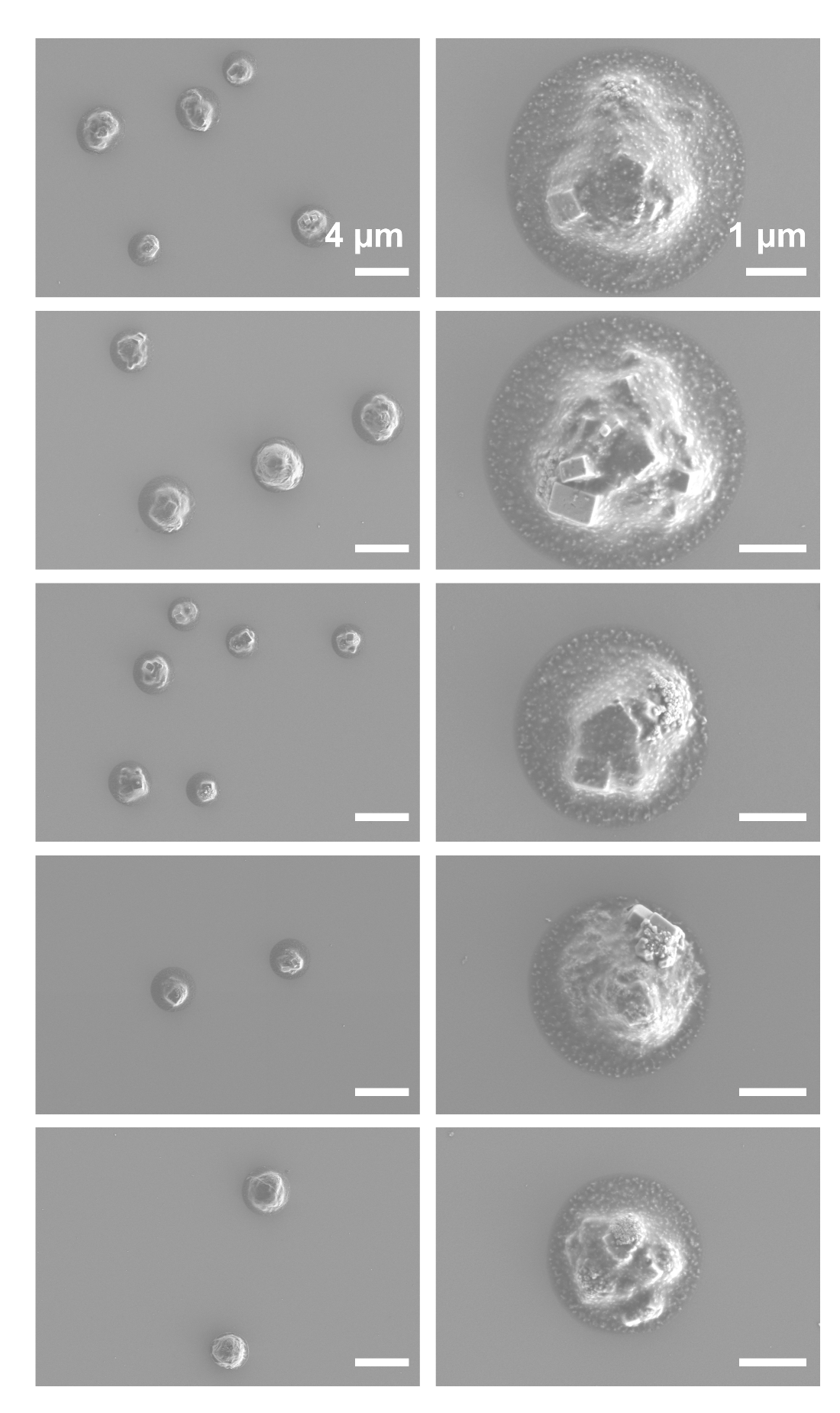

**Figure S20. SEM images for AS–W aerosols**

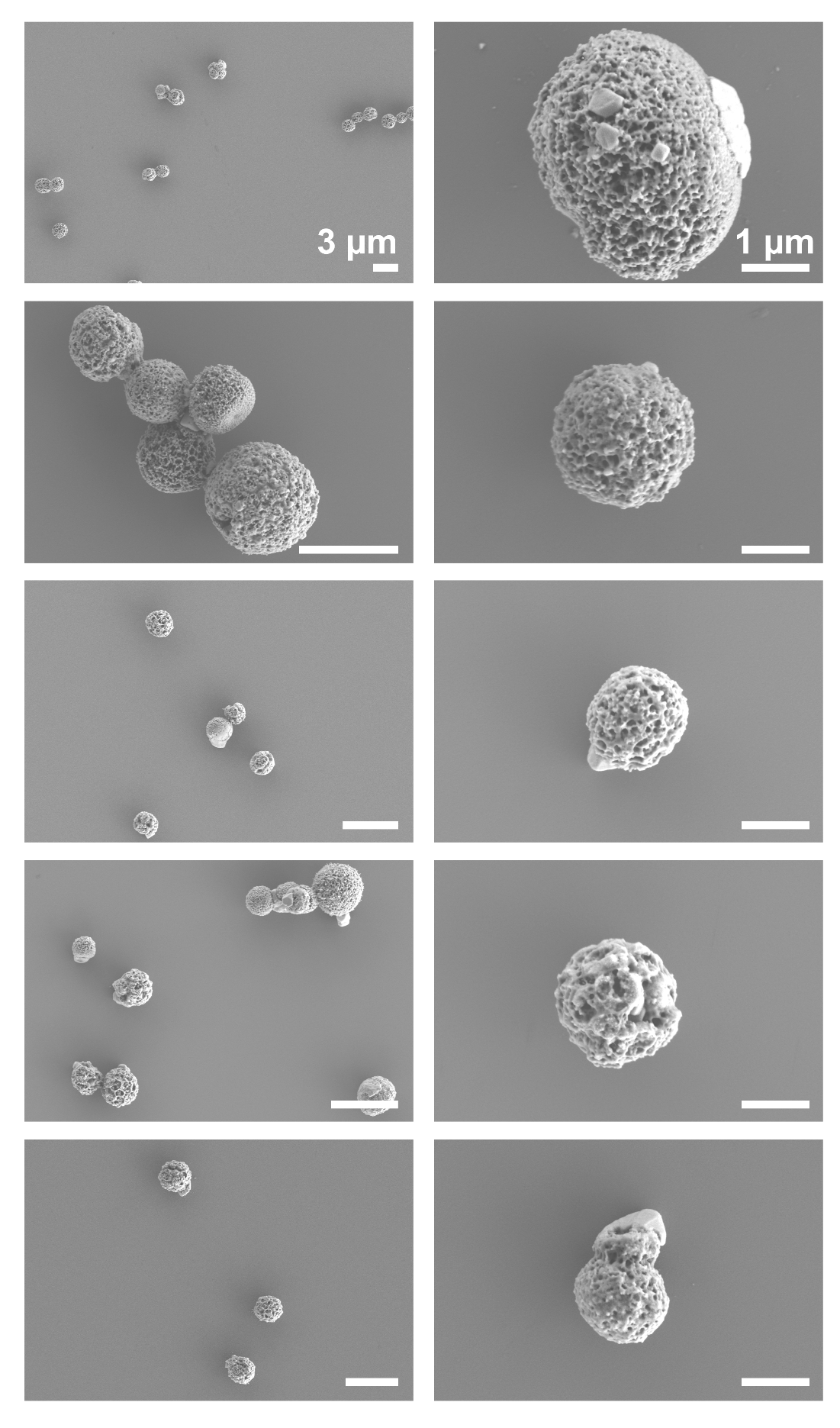

**Figure S21. SEM images for ALF aerosols**

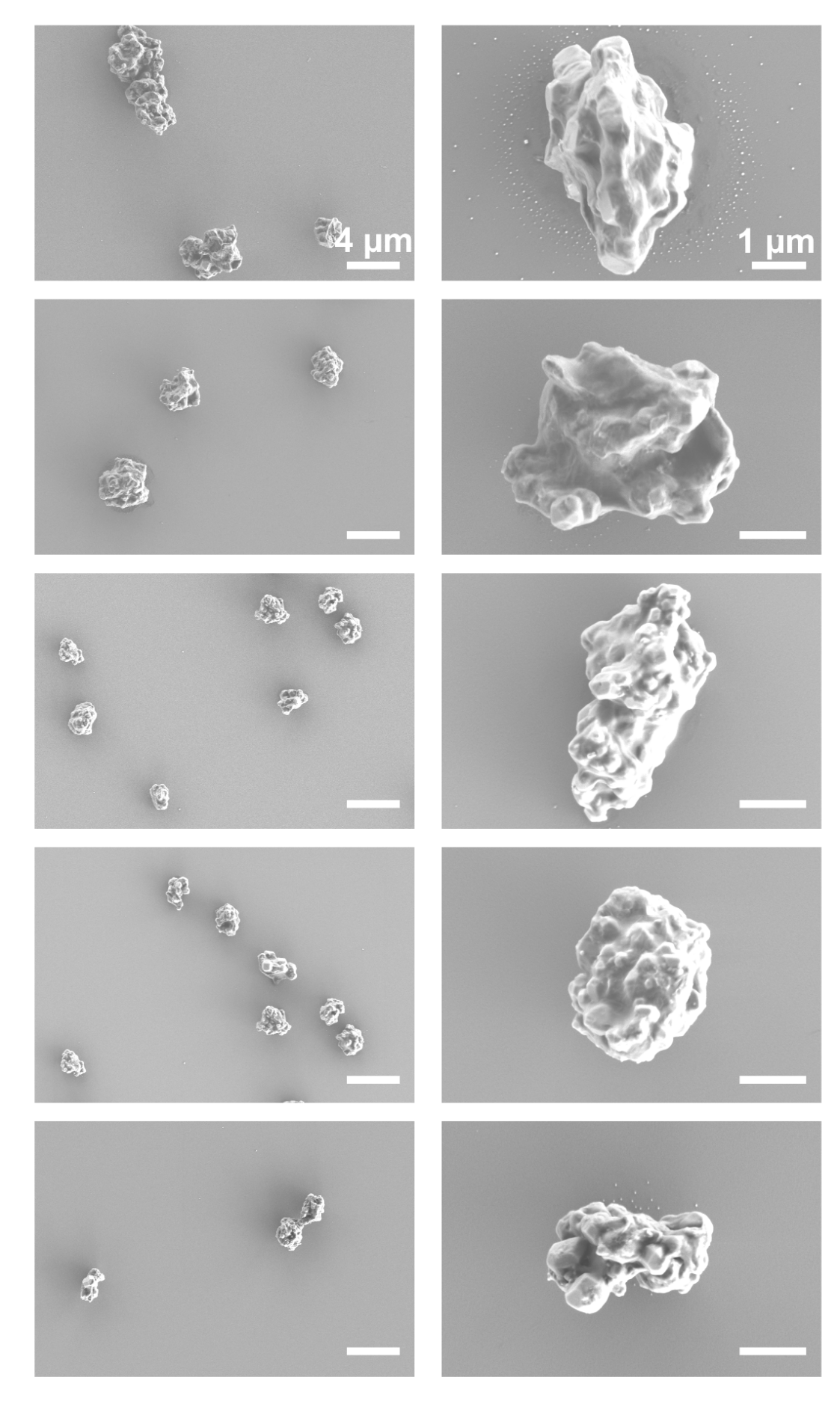

**Figure S22. SEM images for ALF–DPPC aerosol**

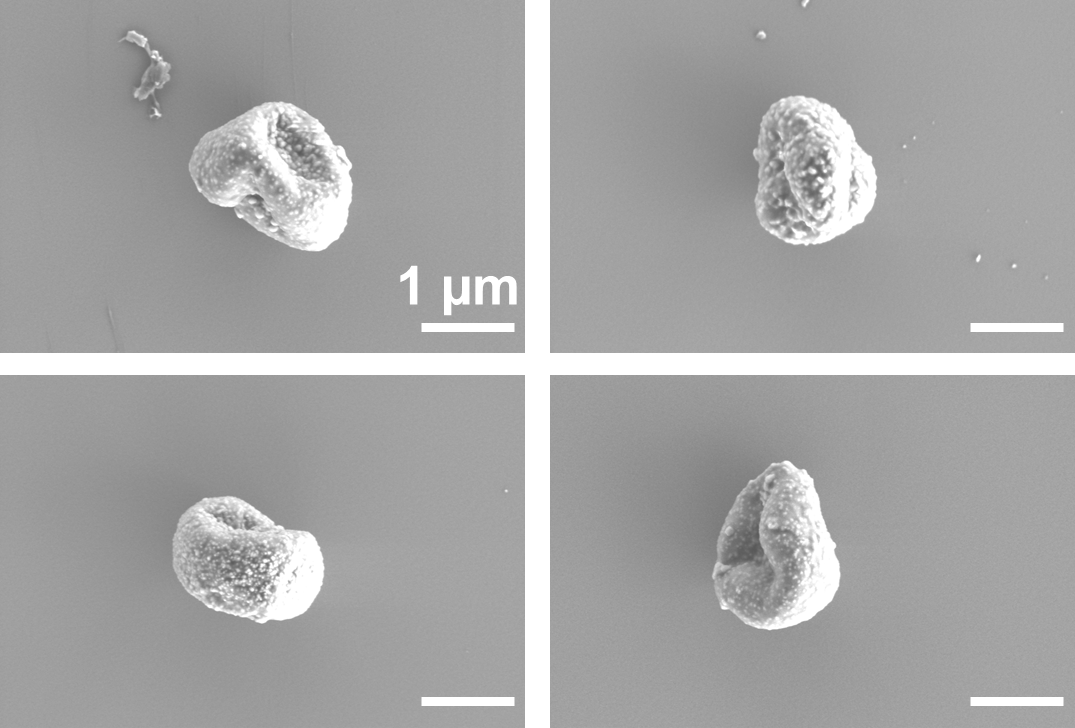

**Figure S23. SEM images of AS–W aerosol particles collected after a residence time of ~50 s**

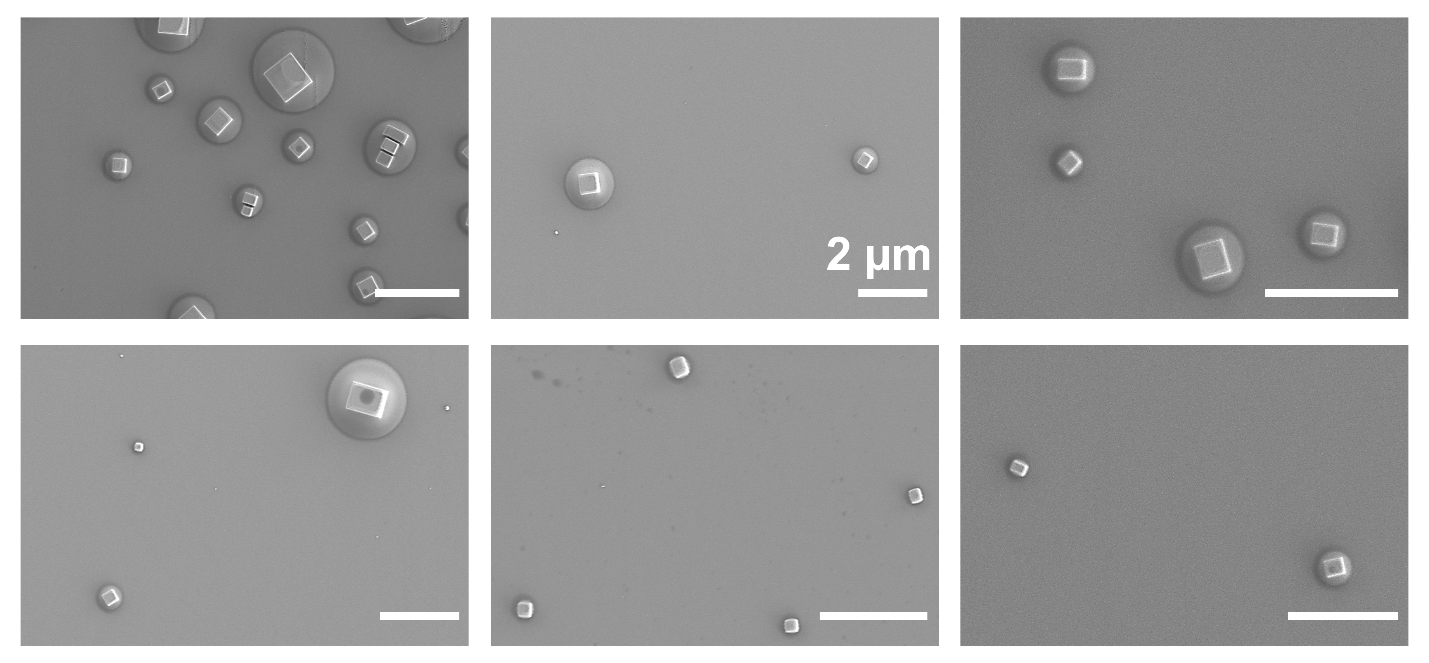

**Figure S24. SEM images of sub-micron NaCl–glucose aerosol particles collected from the major flow of the Virtual Impactor.**
